# Polymer overlap and the organization of enzyme activity in crowded solutions

**DOI:** 10.64898/2026.11.19.767803

**Authors:** Josipa Cecic Vidos, Giuliano Migliorini, Josef Hamacek, Francesco Piazza

**Author notes:** J.C.V. and G.M. contributed equally to this work as co-first authors.

## Abstract

Macromolecular crowding shapes biochemical reactions in subtle ways, yet a general framework connecting enzyme activity to the physical properties of polymeric crowders remains elusive. Here we show that polymer scaling concepts provide a common frame-work to organize the effects of crowding on enzyme activity. Using dextran as a model crowder and fluorescence measurements extending beyond the initial-rate regime, we measure the time course of product formation upon peptide hydrolysis by a representative protease. Time-dependent analysis with a reversible kinetic model yields an inferred equilibrium constant whose dependence on dextran concentration collapses onto a common exponential curve after rescaling by a molecular-weight-dependent concentration. We show that the latter displays the same scaling law as the overlap concentration expected for branched dextran, linking quantitatively the reaction response to a fundamental property of polymer solutions. The common behaviour breaks down for the largest dextran at concentrations well above this scale, identifying the limits of the common scaling behaviour. Altogether, these findings suggest that polymer overlap is a general organizing principle for interpreting biochemical kinetics in polymer-crowded solutions.

## 1. Introduction

Macromolecular crowding influences biochemical reactions through several competing effects, making its overall impact on enzyme activity difficult to predict [1, 2]. Crowders can increase solution viscosity and hinder molecular self-diffusion [1], while interactions with the surrounding medium can stabilize or destabilize enzyme conformations [3–5], affecting the equilibrium of biochemical reactions [6, 7]. Diffusion-limited reactions are expected to slow down as diffusivity decreases [8], whereas crowding can favour activated processes when it stabilizes the transition state relative to the reactants [1, 8]. The cumulative effect therefore depends on the specific features of both the reacting molecules and the crowding agents. For example, stabilization of compact enzyme conformations has been associated with increased activity [9]. For α-chymotrypsin, addition of PEG was reported to decrease both *K_m_* and k*_cat_* through hydration effects [4], whereas dextran increased *K_m_* while decreasing *k_cat_* [10].

A central question is whether some of this diversity can be understood through physical properties shared by different crowding conditions. Synthetic polymers are widely used as model crowders, yet their polymeric nature is not always explicitly incorporated into the interpretation of biochemical assays [11, 12]. Flexible polymers can produce effects that differ from those of globular proteins [13, 14], and their molecular weight, architecture and interactions with the solvent determine how their properties change with concentration [15, 16]. Polymer physics provides scaling concepts that connect these features and may therefore offer a common basis for interpreting their effects on biochemical reactions [15–17].

One such concept is the overlap concentration, *c^∗^*, which marks the crossover from isolated polymer coils to overlapping chains. For polymers whose characteristic size scales with molecular weight as *R ∼ M ^ν^*, the overlap concentration scales as *c^∗^ ∼ M/R*^3^ *∼ M* ^1−3^*^ν^* [15–17]. The reduced concentration *c/c^∗^* thus compares solutions of different polymer molecular weights relative to the onset of overlap, rather than at the same mass concentration alone. This suggests a testable possibility: if the effects of polymer crowding on a biochemical reaction, including shifts of chemical equilibria [6, 7], reflect the same physical properties that organize polymer solution behaviour, reaction responses measured at different molecular weights may admit a common description in terms of *c/c^∗^*.

Testing this possibility also requires following the reaction beyond its initial-rate regime. A recurrent practice in crowding studies is to analyse enzyme activity using initial-rate measurements and the conventional tools of enzymology, whose limits of applicability have already been discussed [18]. Such measurements characterize the early reaction dynamics but do not by themselves determine how crowding affects product accumulation or the long-time balance between substrate and products. Here, we examine progression curves over extended times and analyse them using a reversible kinetic model, seeking information about the asymptotic state of the reaction. This approach allows us to investigate whether parameters inferred from the full progression curves exhibit a common dependence on polymer concentration and molecular weight.

Dextran provides a suitable model system for this purpose. It is a branched polysaccha-ride that has been widely studied and is routinely used in medicine, pharmacology and the food industry [19, 20], as well as in crowding experiments [10, 12]. Its degree of branching depends on its biological origin and synthesis conditions [20], with typical values of around 5% for commonly used dextran samples [21]. Reported size exponents of approximately ν *≃* 0.44 are consistent with its branched architecture [21, 22]. Although hyperbranched and randomly branched polymer models have been considered, its assignment to a specific class of branched polymers remains unresolved [23]. In a recent study, we combined complementary experimental techniques and modelling to characterize the physical properties of dextran solutions within a common polymer-physics framework [24]. This characterization provides a physical basis for examining whether the same scaling concepts can organize the effects of dextran crowding on enzyme activity.

As a model reaction, we investigate peptide hydrolysis by human neutrophil elastase (HNE), a member of the chymotrypsin-like serine protease family [25, 26]. HNE is released by activated neutrophils and acts in the extracellular matrix, where it contributes to tissue remodelling and the regulation of inflammation [27, 28]. Its activity has generally been characterized in standard buffers [29–31], although its extracellular environment is complex and crowded. The catalytic mechanism of serine proteases is well established [26]: substrate binding is followed by enzyme acylation and release of a first product, then deacylation and release of a second product. HNE therefore provides a well-characterized enzymatic system in which to examine how polymer crowding modifies reaction progression.

In this work, we combine fluorescence measurements with kinetic modelling to investigate HNE activity up to long times in solutions containing dextran of three molecular weights. A calibration method recently developed by our group allows us to follow product formation at high concentrations of fluorophores, substrates and crowders [32]. We compare conventional initial-rate analysis with simultaneous fits of extended progression curves using a kinetic model that includes reverse reactions. Dextran increases product formation across the conditions investigated, while the equilibrium constant inferred from the model shows a molecular-weight-dependent concentration response. These responses collapse onto a common exponential curve when concentration is rescaled by a characteristic concentration whose molecular-weight dependence is consistent with dextran overlap and its branched architecture [24]. Deviations emerge for the largest dextran at high concentrations.

Together, our results support the conjecture that polymer overlap can provide an organizing principle for interpreting biochemical reactions in synthetic polymer-crowded solutions and define a basis for testing this description across other biochemical reaction systems and polymer architectures.

## 2. Materials and methods

### 2.1. Chemicals and Starting Materials

7-Amino-4-methylcoumarin (AMC) and hydrochloric acid (p. a., 37 %) were purchased from Fisher Scientific. The fluorescent substrate for elastase (M-AMC: MeoSuc-Ala-Ala-Pro-Val-AMC, elastase substrate V, Calbiochem), Dulbecco’s Phosphate Buffered Saline (10x concentrated, PBS) and Human Neutrophile Elastase (HNE, CAS 9004-06-2, Calbiochem) were acquired from Sigma, as well as three dextrans from *Leuconostoc sp.* : DEX10 (MW range 9 000 to 11 000 Da, average *M_w_* = 9126 Da), DEX40 (35000 - 47000 Da, average *M_w_* = 36720 Da) and DEX450 (450 000 to 600 000 Da, average *M_w_* = 476350 Da). Average values and ranges of *M_w_* are declared by the manufacturer. Dimethyl sulfoxide (DMSO, 99 % purity, for molecular biology) was obtained from Acros Organic. The degradation product of the elastase substrate, a small peptide (M : MeoSuc-Ala-Ala-Pro-Val) was synthesized by the SynBio3 platform at the Institute of Biomolecules Max Mousseron (UMR5247 CNRS-ENSCM-University of Montpellier) and used without further purification.

### 2.2. Stock Solutions

Solutions of 7-amino-4-methylcoumarin (AMC) and elastase substrate were prepared by dissolving the powders in DMSO at a concentration of 15 mM. The preparation of stock solutions involved subsequent dilutions using 10x Dulbecco’s Phosphate Buffered Saline (PBS) to have a final concentration of 1x PBS.

The HNE was reconstituted by incubation for 20 minutes at room temperature followed by the addition of 200 µL of the reconstitution buffer (50 mM sodium acetate, 200 mM NaCl, 0.1% BSA, pH 5.5) and incubated for 20 min with gentle shaking at 22°C. The stock solution of the enzyme had the concentration of 16.95 µM and was aliquoted and stored at *−*20°C until use. Before using the enzyme for the reactions, enzyme solutions were diluted to 450 nM using 10x PBS and MiliQ water to reach pH 7.4. The final concentration of enzyme during the reactions was 45 nM.

All solutions were used immediately or stored at *−*20°C until required, ensuring optimal freshness and stability for experimental use except for dextran solutions kept at +4°C until use. Stock solutions of dextrans 10, 40 and 450 kDa (in the following DEX10, DEX40 and DEX450) were prepared by dissolving an appropriate amount of powder in 10x PBS and water. The solutions were then homogenised using a Homogeniser 850 (Fisher Scientific, USA) using 7 mm × 115 mm dispersion element (11000 rpm, 7 cycles, each lasting 3 min with a 1 min break). Stock solutions were diluted with PBS to the intermediate concentrations used in the experiments to obtain final concentrations of 5, 10, 15 and 20 % (w/w). The weight fraction is obtained by measuring the mass of crowder before adding the solvent and dividing it by the mass of solution w = *M_c_*/*M_tot_*. In section 6 dextran concentrations are referred to as mass concentrations *c* = *wρ(w)*, where ρ*(w)* is the density of dextran solutions at 37°C. The latter were directly estimated with a METTLER-TOLEDO Density 30 px densitometer, and fitted to the relation

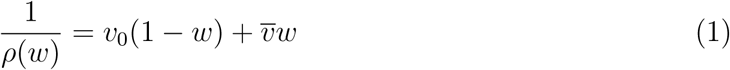

obtaining v_0_ = 1/ρ_0_ = (0.9972 *±* 0.0011) cm^3^/g and *v̅ ≈* 0.63 cm^3^/g. The measurement of density and the discussion of its implications have already been presented in another manuscript [24].

### 2.3. Measurement of HNE kinetics

The catalytic action of Human Neutrophile Elastase was assayed using the substrate MeoSucc-Ala-Ala-Pro-Val-AMC, elastase substrate V (Calbiochem), which contains a fluorophore 7-amino-4-methyl coumarin (AMC). AMC is not fluorescent when covalently bound to the peptide, but its fluorescence is recovered after enzymatic cleavage, with excitation at 370 nm and emission at 450 nm. The reactions were followed at 37° C for 2 hours. Samples containing the working buffer, HNE and dextran (total volume of 90 µl) were prepared in black-bottomed 96-well plates (Nunc™ MicroWell™ 96-Well, Nunclon Delta-Treated, Flat-Bottom Microplate). Plates were pre-incubated for 10 minutes inside the plate reader ClarioStar (BMG Labtech). The automatic injection was utilised to start the reaction by injecting 10 µL of the substrate of appropriate concentration into the wells to keep the final volume of 100 µL. After the injection of the substrate, the plate was shaked for 180 seconds at 300 rpm to ensure homogeneous mixing. To prevent evaporation, the plate was securely sealed with a clear adhesive film (VWR, adhesive film for PCR plates, polyester, 50 µm ultra-clear film). Fluorescence intensity was then measured for 2 hours with cycles of 90 seconds to follow the progression of the reaction. The substrate concentrations used were 0, 80, 150, 300 and 750 µM of substrate. Each plate contained a range of concentrations of the AMC fluorophore and the control reaction with 150 µM substrate used to verify the repeatability between the plates.

### 2.4. Fluorescence spectra and calibration curves

In order to convert fluorescence intensity units (FI) to the concentration of the product generated by the enzyme, calibration curves were measured from separate experiments performed in the same conditions as HNE kinetics. In short, solutions of AMC were prepared, incubated inside the plate reader at 37° C and covered with clear adhesive films. Plates were measured for 1 or 2 hours until the loss of fluorescence over time was no longer significant. Concentrations of AMC comprised the range 0, 2, 6, 15, 50, 150, 400, 750 µM. To investigate the potential impact of substrates and dextran on the fluorescence intensity of the product, supplementary sets of samples were prepared. In the presence of AMC (0, 2, 6, 15, 50, 150, 400, 750 µM) two different concentrations of substrate (80 and 750 µM) and two concentrations of dextran (10 and 20% (w/w) of each size) were introduced. The procedure of conversion of FI to the concentration of the product and a discussion of the influence of dextran and substrate is elaborated in Section 3.

### 2.5. Fit methods

In order to fit kinetic models of enzymatic catalysis to our time series, we solved numerically the systems of ordinary differential equations (ODEs) for the time-dependent concentrations 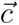 of the form

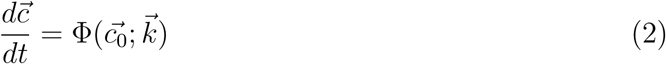

The function Φ is completely characterised by the initial concentrations 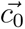. The substrate (s_0_) and enzyme (e_0_ = 45 nM) initial concentrations were fixed by the experimental conditions. Initial concentrations of product and intermediate species were set to zero. The vector 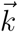 comprises the rate constants that characterise the kinetic model. The length of both vectors 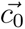 and 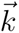, as well as the number of equations in system Φ, depend on the choice of the specific kinetic scheme adopted.

The numerical solutions of the system (2), 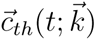, characterised by 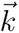, were optimized by minimizing the error-weighted distance between experimental product concentrations, *P_n_*, and the model predictions, evaluated at the same times *t_n_*. ODE systems representing chemical reactions can be stiff if kinetic constants differ by several orders of magnitude. This problem was taken care of using the LSODA method [33] in the scipy.integrate package in Python, which enabled us to handle the stiffness encountered in the fitting procedure, without significant extra computational cost.

The fits were performed using an original global approach. Time series corresponding to different initial substrate concentrations s_0_ were fitted using a single set of kinetic parameters. In practice, we minimized the following reduced chi square function

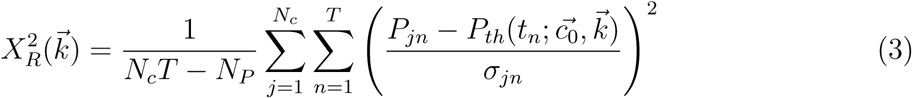

Here, *N_c_* = 4 is the number of substrate concentrations, T is the number of time points, *N_P_* is the number of kinetic constants, while *P_jn_* and σ*_jn_* are the product concentrations and the corresponding errors measured at time *t_n_* in the experiment performed with the j-th initial substrate concentration. The minimization was realized through the combination of a home-made stochastic optimization algorithm, reminding of a Metropolis-Hastings strategy at finite temperature, where the cost function 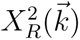 plays the role of the energy, coupled to a final steepest-descent refinement. To estimate the uncertainties associated with the best-fit values of the free parameters, the covariance matrix was obtained by sampling around the minimum of the cost function with our stochastic algorithm.

## 3. Fluorescence calibration

Observing enzymatic kinetics through the release of a fluorescent product in a multicomponent mixture can be challenging. While in dilute conditions fluorescence is proportional to fluorophore concentration, this is no longer the case at high concentrations. For example, high concentrations of substrates can lead to absorption mechanisms, mainly by the released fluorophores, through the so-called inner filter effects [34].

Comparing kinetics in different crowding conditions poses an additional challenge. At high concentration of macromolecules [35], optical mechanisms involving the crowders them-selves cannot be neglected. Consequently, a preliminary careful characterization of the optical properties of the mixture is required in order to obtain a reliable calibration. Here, we follow a previously described strategy to calibrate a spectrofluorometer with front-face geometrical setup, in the presence of complex mixtures of chemical species [32]. The fluorescence intensity emitted by a solution composed of n species at concentrations *c_i_*, as long as scattering can be neglected, is described by the following relation

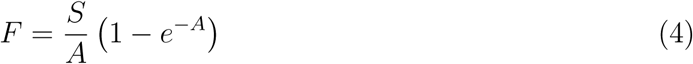

with

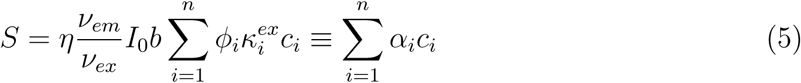

and

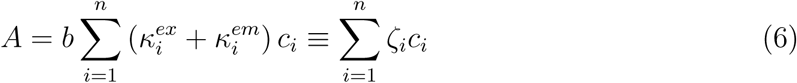

The source term S accounts for emission processes by the species in solution. Overall, the parameters α*_i_* comprise the contributions from the molar absorption coefficient 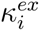 at the excitation wavelength, fluorescence quantum yield ϕ*_i_*, excitation intensity *I*_0_, emission and excitation frequencies *ν_em_* and *ν_ex_*, and optical length of the sample b. The factor η accounts for the geometry of the experimental setup and for the conversion to dimensionless units. A is the absorbance of the sample, expressed as a linear combination of the concentrations *c_i_* weighted by their molar absorption coefficients 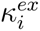 and 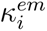 at excitation and emission wavelengths, respectively. Further details on the derivation of relations (4), (5) and (6) are available from our recent paper [32]. At high concentrations, one expects the contributions to absorbance by any species in solution. As a consequence, the calibration relation is expected to be a non-linear function of several concentrations, reducing to the usual linear relation in the dilute limit.

### 3.1. Experimental determination of the calibration relation

In order to characterise S and A of a complex mixture containing fluorophores, substrates and crowders, separate calibrations of binary mixtures containing the fluorophore and either the substrate or a crowder were conducted. The results of the calibration in the presence of substrate and DEX10 are listed in Table 1 and illustrated in Figures S1 and S2. Interestingly, at the wavelengths of interest, only absorption by DEX10 turned out to be relevant. Extinction by DEX40 was not observed, while that from DEX450, even though detectable, also could not be resolved and was therefore neglected (Figures S3 and S4). Absorption spectra were collected too, to confirm the results of fluorescence calibration and to check that scattering could be neglected [32].

**Table 1:**
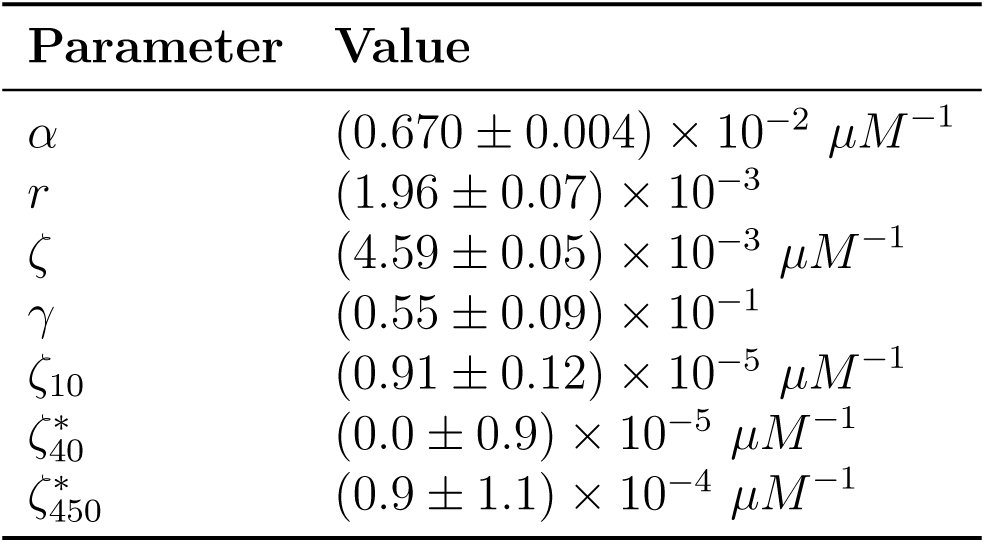
Best estimates of the calibration parameters obtained for different binary mixtures with product (AMC) and substrate (AMC-peptide) or dextrans. α, r, ζ and γ were chosen as the best-fit result from the calibration in the presence of substrate (Figures S1 and S2). ζ_10_ was estimated from the calibration relation in the presence of DEX10, keeping α and ζ fixed to the values obtained in the absence of crowders. The same fitting procedure was followed for DEX40 and DEX450. For the former, no absorption was detected, while for the latter it was compatible with zero and was therefore neglected. Therefore, no correction for crowder absorption was adopted for these two sets of data. Additional information about the calibration relation is reported in the S.I..

| **Parameter** | **Value** |
|---|---|
| α | (0.670 ± 0.004) × 10<sup>-2</sup> µM<sup>-1</sup> |
| *r* | (1.96 ± 0.07) × 10<sup>-3</sup> |
| ζ | (4.59 ± 0.05) × 10<sup>-3</sup> µM<sup>-1</sup> |
| γ | (0.55 ± 0.09) × 10<sup>-1</sup> |
| ζ<sub>10</sub> | (0.91 ± 0.12) × 10<sup>-5</sup> µM<sup>-1</sup> |
| ζ<sub>40</sub><sup>*</sup> | (0.0 ± 0.9) × 10<sup>-5</sup> µM<sup>-1</sup> |
| ζ<sub>450</sub><sup>*</sup> | (0.9 ± 1.1) × 10<sup>-4</sup> µM<sup>-1</sup> |

Overall, the result of our calibration allowed us to estimate the fluorescence intensity of the mixture through the relation (4), with

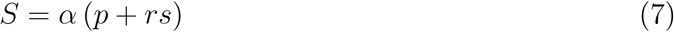

and

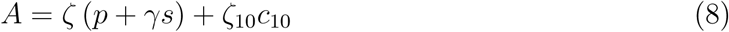

where *p*, *s* and *c*_10_ are the molar concentrations of fluorophore (product), substrate and DEX10 ^∗^, respectively, α and ζ are the dominating contributions that come from fluorophores, while the factors r, γ and ζ_10_ are corrections due to the other species in solution. We refer to Table 1 for the best estimates that were used to treat the data collected in kinetic experiments.

Finally, *F* must be converted into a molar product concentration *p*. In general, during the enzymatic reaction, both *s* and p depend on time. Assuming that the initial product concentration *p*_0_ is exactly zero, then the sum of *s* and *p*, at any time during the reaction, lies in the narrow interval *s*_0_ *− *e**_0_ *≤ *s** + *p ≤ s*_0_ (*s*_0_ *− (*s + p*) ≤ *e**_0_ as *e*_0_ is the upper limit of the concentration of the intermediate states). In order to invert relation (4), we approximate the substrate concentration as s *≃ s*_0_ *− p − e*_0_/2, so that s + p takes the mean value of the interval. With substrate and product concentrations in the 10 - 1000 µM range and e_0_ = 45 nM, this approximation is amply justified. The relation (4) becomes a monotonic function of *p*,

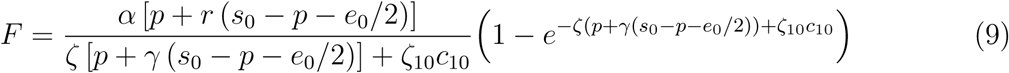

Therefore, one and only one value *p_n_* can be associated with each point of the measured fluorescence time series *F_n_* by solving numerically the equation *F_n_ − F*(*p_n_*) = 0. We used standard error propagation techniques to compute uncertainties at each time point, σ*_n_*, which include the experimental errors on *F_n_* and the contribution from the fitted parameters in Table 1 (more information can be found in the S.I.).

## 4. Short-time analysis of the enzyme kinetics

### Enzymatic kinetics in diluted solutions

Usually, an enzymatic activity is characterised through the Michaelis-Menten (MM) affinity *K_m_* and the maximum velocity, *V_M_* = *k_cat_*e_0_, estimated from a MM plot of the initial slopes of the product time series. Consensus values of these parameters for the degradation of M-AMC and M-pNa by HNE are reported in Table 2. These identify a range to compare our results with, even though our initial velocities are recorded after roughly 300 s from the start of the reaction. Moreover, the experimental conditions could be different from ours for some of these studies.

**Table 2:** Consensus analysis of the kinetic parameters for the hydrolysis of substrates MeoSuc-AAPV-X (M-X) by Human Neutrophil Elastase and Suc-AAPV-AMC (S-AMC). k_2_ and k_3_, the acylation and deacylation kinetic rates, are reported from studies by Stein and collaborators who considered pre-steady-state burst amplitude data in a sub-second timescale [30]. In these studies, *K_s_* = (*k_off_* + *k*_2_) /*k_on_*, is different from *K_m_* = (*k_off_* + *k_cat_*) /*k_on_*. In ref. [31] k_3_ is assumed to equal *k_cat_* for the hydrolysis of the corresponding thiobenzyl ester: MeoSuc-AAPV-SBzl (M-SBzl); k_2_ and *K_s_* are determined consequently. All other kinetic constants are estimated from fits of steady state initial rate velocities, with the Lineweaver-Burk method, fit to the Michaelis-Menten equation, or other standard methods.

| Subs. | Enz.<br>conc.<br>[*µM*] | Subs.<br>conc.<br>range<br>[*µM*] | *K<sub>m</sub>*<br>[*µM*] | *k<sub>cat</sub>*<br>[*s*<sup>−1</sup>] | *k<sub>cat</sub>*/*K<sub>m</sub>*<br>[*M*<sup>−1</sup>*s*<sup>−1</sup>] | *k*<sub>2</sub><br>[*s*<sup>−1</sup>] | *k*<sub>3</sub><br>[*s*<sup>−1</sup>] | *K<sub>s</sub>*<br>[*µM*] | Exp. cond. | Ref. |
|---|---|---|---|---|---|---|---|---|---|---|
| M-AMC | 1.8 × 10<sup>−2</sup> | 89 - 200 | 290 | 3.3 | 1.1 × 10<sup>4</sup> |  |  |  | pH = 7.5; 0.05 M THAM buffer, 0.5 M NaCl, 0.01 M CaCl, 9.9% v/v DMSO; T = 25°C. | [36] |
| M-AMC |  |  | 138 ± 16 | 16.8 ± 1.2 | (1.06 ± 0.09) × 10<sup>5</sup> |  |  |  | pH = 7.4; 0.010 M Hepes buffer, 140 mM NaCl and 0,05% Triton X-100; T = 37 °C. | [37] |
| M-AMC | 0.100 |  | 923 ± 52 |  |  |  |  |  | buffer details : [38] ; T = 37 °C. | [38] |
| M-pNA | 0.100 |  | 271 ± 38 |  |  |  |  |  | buffer details : [38] ; T = 37 °C. | [38] |
| M-pNA |  | 50 - 4900 | 140 | 17 | 1.2 × 10<sup>5</sup> |  |  |  | pH = 7.5, ; 0.1 M Hepes buffer, 0.5 M NaCl, 9.8% dimethylsulfoxide ; T = 25°C. | [39] |
| M-pNA |  | 20.5 - 410 | 54 ± 4 | 10.0 ± 0.2 | (1.85 ± 0.14) × 10<sup>5</sup> |  |  |  | pH = 7.37 ; 0.100 M phosphate, 0.50 M NaCl, 3.3% *Me*<sub>2</sub>*SO* ; T = 25.0 ± 0.1 °C. | [29] |
| M-pNA | 42 |  | 55 ± 6 | 13.3 ± 0.8 | 2.4 × 10<sup>5</sup> | 30 ± 4 | 23 ± 6 | 115 ± 7 | pH = 7.4, ; 0.100 M phosphate, 0.500 M NaCl,0.020 M acetate, 5% *Me*<sub>2</sub>*SO* ; T = 29 °C. | [30] |
| M-pNA |  |  | 55 | 10 | 1.82 × 10<sup>5</sup> | 43 | 13 | 240 ± 7 | pH = 7.4 ; 0.100 M phosphate, 0.5 M NaCl, 3.3% *Me*<sub>2</sub>*SO* ; T = 25.0 ± 0.1 °C. | [31] |
| M-pNA |  | 280 - 1000 | 210 | 23 | 1.12 × 10<sup>5</sup> |  |  |  | pH = 7.4 ; 0.050 M Tris/HCl 0.1OO M NaCl, and 0.05% Triton X-100 ; T = 37 °C. | [40] |
| S-AMC |  | 15.6 - 1000 | 260 | 21 | 0.82 × 10<sup>5</sup> |  |  |  | pH = 7.6, ; 0.1 M NaH2PO4, 0.5 M NaCl, ; T = 37 °C. | [41] |

To estimate short-time reaction velocities, we fitted a straight line to the first five points of the product time series, *P*(t) = *v*(*t − t^∗^*). The time *t^∗^* is seen to be negative for all concentrations. This means that the velocities that we measure from the first points are lower than the average velocity during the blind window. Consequently, the real initial velocities *dP* (0)/*dt* are higher than *v*.

A standard analysis would proceed by building a MM plot based on v vs s_0_ and determine the maximum velocity *V_M_* and the Michaelis constant *K_m_*through the usual relation

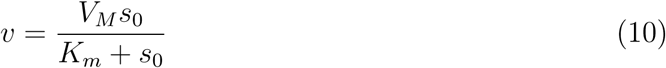

The fit (Figure 1) gives *V_M_* = (0.145 *±* 0.006) µM/s and *K_m_* = (1.23 *±* 0.08) mM. From the value of *V_M_* and *e*_0_ = 45 nM, we readily get *k_cat_* = *V_M_* /*e*_0_ = (3.2 *±* 0.1) *s^−^*^1^.

**Figure 1:**
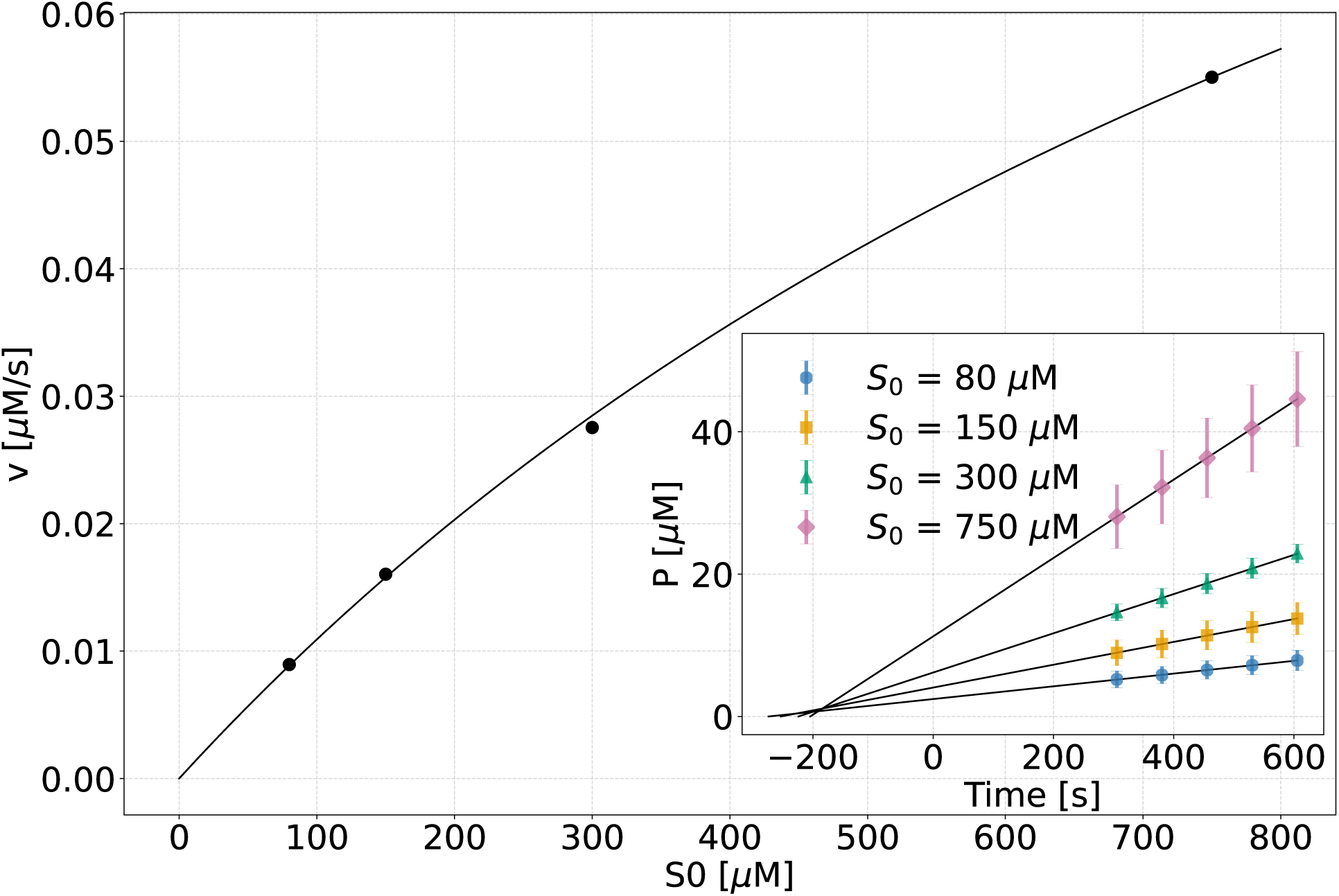
Michaelis Menten plot. Velocities are estimated from linear regressions to the first five points of the product progress curves (inset).

The *K_m_* value is close to that reported by ref. [38], but 5-10 times higher than those reported in ref. [36] and [37], while *k_cat_* is close to the original estimation found in ref. [36] at T = 25 °C, and lower than the one reported by [37] at our working temperature T = 37 °C. Overall, a blind MM analysis such as this tends to underestimate the maximum velocity. Based on this discussion, however, we expect that an extended-time analysis will give the somewhat higher correct values of *V_M_* in agreement with the literature.

Even though two hours may be considered a long duration for an enzymatic fluorometric assay, a direct inspection of Figure 2 shows that this time lapse is not enough to reach the equilibrium state. It is instructive to examine the trend of the reaction velocity *dP*/*dt* as a function of time, estimated as simple finite difference of the product time series (Fig. S5). It is apparent that the reaction velocity is never stationary in our experimental time range.

**Figure 2:**
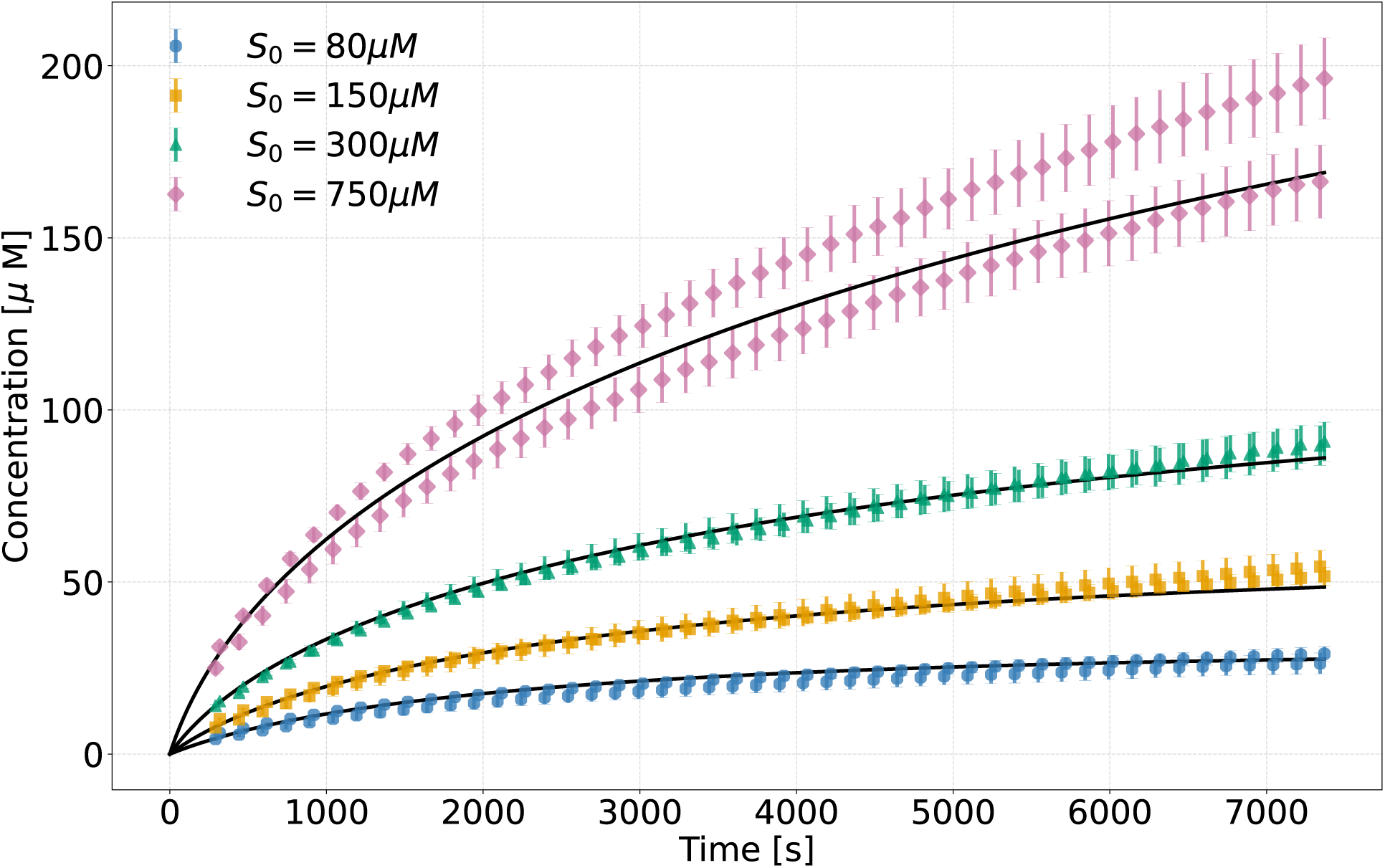
Measured evolution of the product concentration in the absence of dextran (symbols) and best-fit with model (12) (solid lines), at different initial substrate concentrations. The data shown and the error bars represent averages and standard errors computed over 4 replicates.

### Enzymatic kinetics in solutions crowded by dextrans

Fig. 4 shows that, also in crowded conditions, one cannot consider that the reaction has reached equilibrium at the end of the observation time range. Nevertheless, we found that the last measured HNE activity increased consistently as a function of dextran weight fraction, for all the investigated molecular weights and concentrations of dextran in solution. Remarkably, we found that the reversible MM scheme (11) also represents an accurate model for the observed kinetics for all substrate concentrations and crowding levels (see fits in Fig. 4), providing a microscopic interpretative framework for the observed crowding-induced acceleration of substrate digestion. This effect is illustrated explicitly in Fig. 3d, which shows that 20 % w/w of dextrans leads to twice as much product at the end of the observation window than in the absence of crowding. Dextran appears then to boost the digestive activity of HNE. However, the short-time analysis tells a different, clearly inaccurate story. The maximum velocity seems to increase with crowding with the notable exception of DEX450, while the affinity shows a consistent decrease with crowding, flagging a stabilization of intermediate enzyme-substrate/product species. Overall the enzyme efficiency moderately increases (Fig. 3c). Overall, our extended-time analysis shows that the usual treatment of short-time catalytic activity fails to thoroughly describe the effect of crowding on the enzyme activity (see again Fig. 3d).

**Figure 3:**
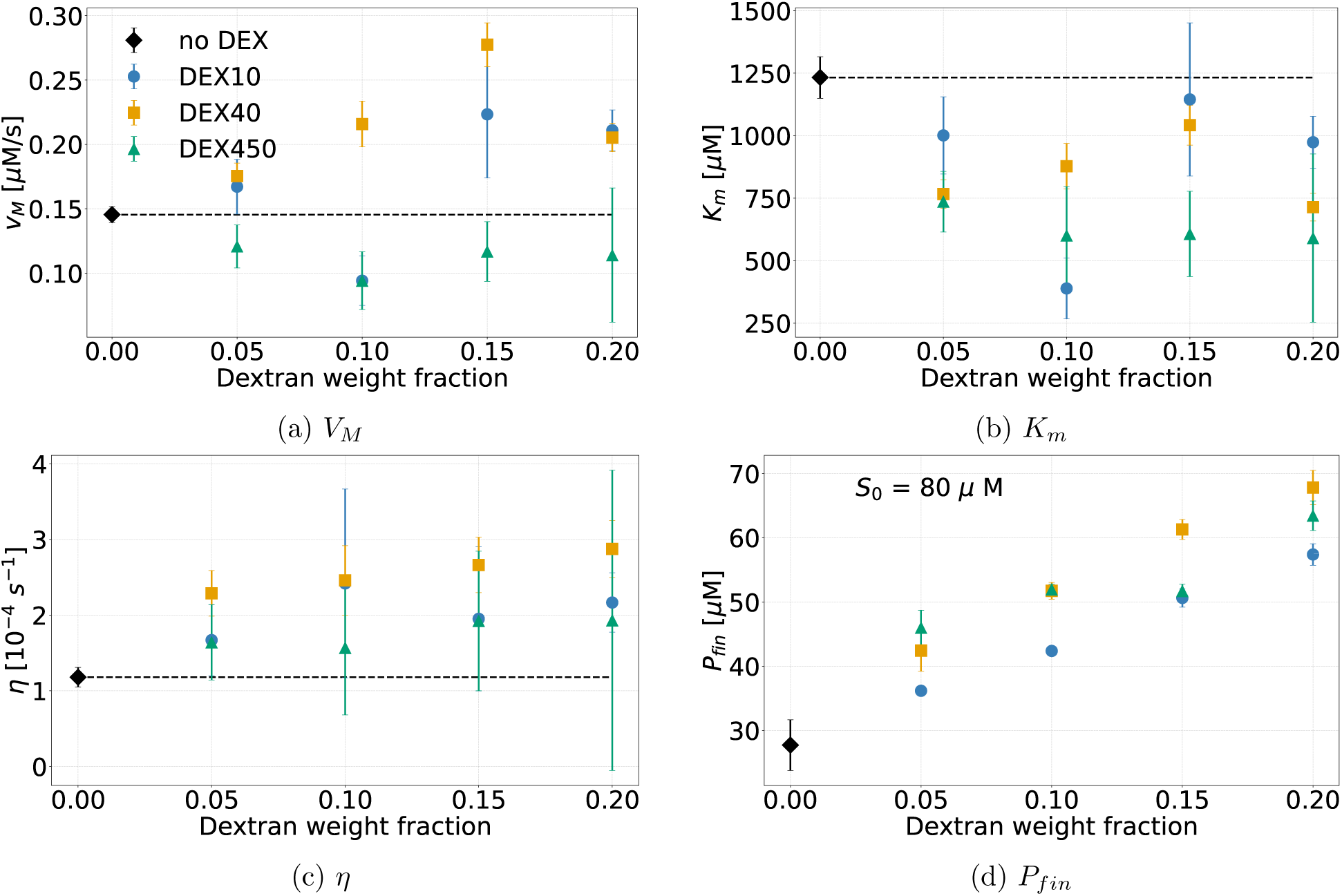
MM parameters extracted from a conventional analysis of the initial slopes compared with long-term product levels. (a) Maximum velocity *V_M_* as a function of dextran weight fraction. (b) Michaelis-Menten constant *K_m_* as a function of dextran weight fraction. (c) Catalytic efficiency *η* = *V_M_ /K_m_* as a function of dextran weight fraction. (d) Comparison of the last measured product levels for *s*_0_ = 80 *µ M*, versus dextran weight fraction.

**Figure 4:**
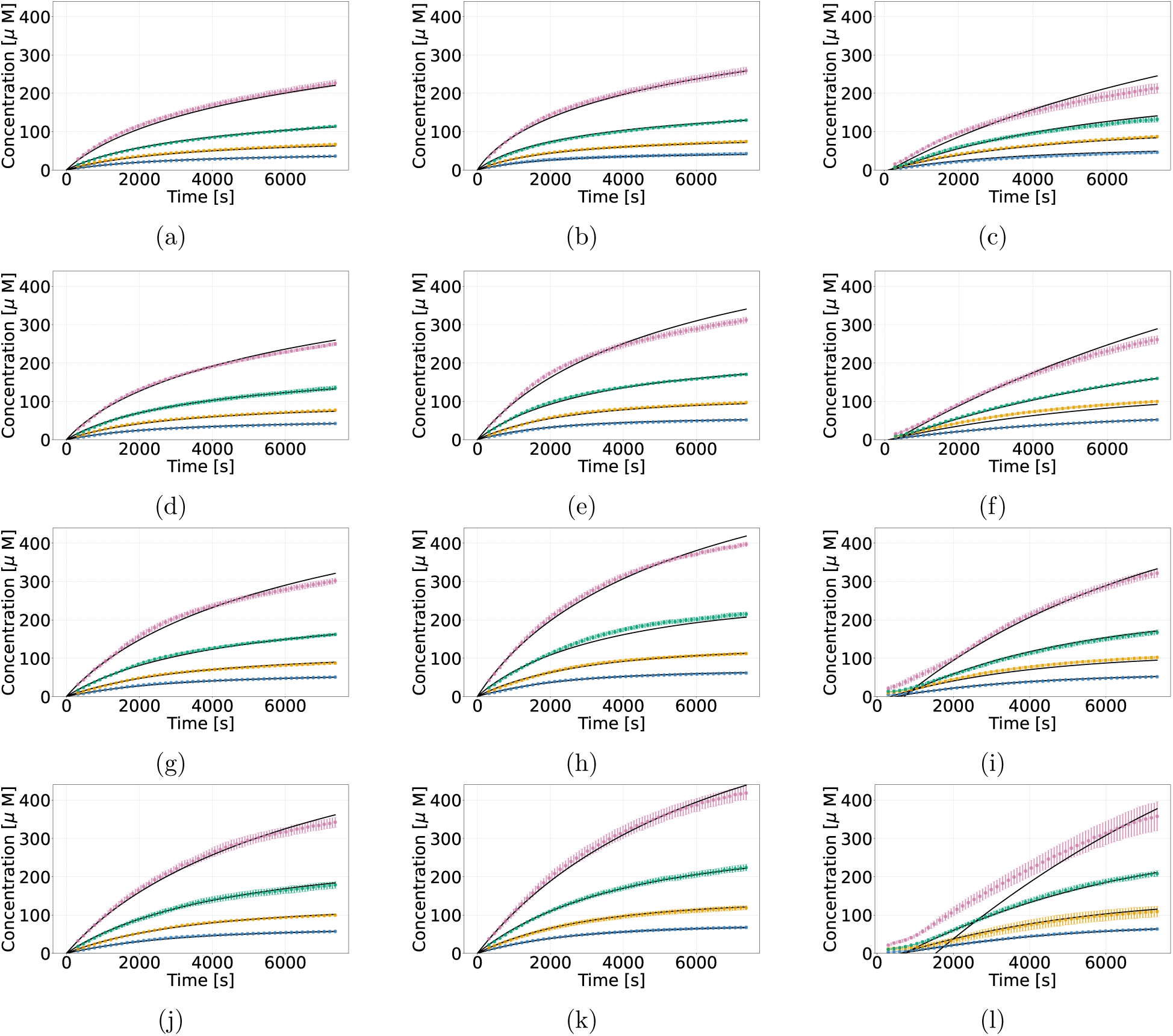
HNE kinetics in the presence of dextran at different weight fractions *w*. Columns: DEX10, DEX40 and DEX450. From top to bottom, dextran weight fractions: 0.05, 0.10, 0.15 and 0.20. DEX450 shows a transient lag time before reaching maximum velocity. Black lines are the global best-fit solutions, obtained by numerically integrating the rate equations corresponding to the scheme (12). In the case of DEX450, the fits neglect a transient, as explained in section 6. The data shown and the error bars represent averages and standard errors computed over 4 replicates.

## 5. Extended-time analysis of enzyme kinetics

Given the manifest incompleteness of the short-time analysis, we sought to develop a method to take into account the extended time series of product generation collected in long experiments. Even though serine proteases are known to degrade their substrates with the three-step mechanism described in the introduction [26], a reasonable starting point is a model with two steps and a single, coarse-grained intermediate complex, X, which dissociates thereby regenerating the free enzyme and releasing the detected product

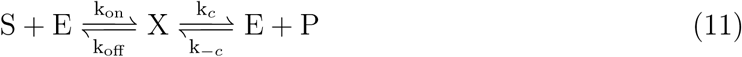

The first step accounts for the association of substrate and enzyme. Contributions from the diffusion of substrates and enzymes, and any further conformational step preceding the formation of X, are accounted for by the kinetics constants in this step, *k_on_* and *k_off_*. In this approach, the second step accounts for both, enzyme acylation, with release in solution of the cleaved AMC fluorophore, and deacylation, which leads to the release of the peptide (M) and the free enzyme in solution. The two steps leading to the consecutive release of products in the serine catalysis are known to occur with kinetic constants whose relative magnitude depends on the nature of the substrate. Literature data in Table 2 suggest that their values might be similar and that they are too fast to be separated in an experiment in the time range of hours. Therefore, the rate constant *k_c_*, accounts for both, enzyme acylation and deacylation. The rate *k_−c_* accounts for recombination of the product(s) with the enzyme, ensuring microscopic reversibility as required by thermodynamic consistency.

Even though the exact nature of this recombination cannot be identified at this level of coarsegraining, considering amino acid -amino acid interactions at the HNE substrate recognition sites [26, 30], it seems reasonable that the peptide M can interact at the active site, saturating elastase activity on the long term. Moreover, since we work in substrate excess, the observed product concentrations are in the micromolar range at any time. Therefore, the concentration difference between the two products, limited by the nanomolar enzyme concentration, is negligible. This justifies the coarse graining of the full kinetic scheme to the single-product mechanism (11). The rate equations read (lower-case letters denote concentrations)

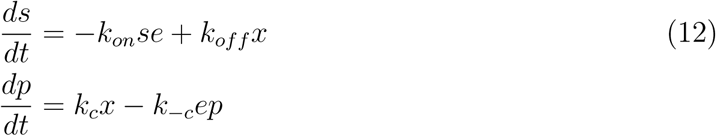

with conservation laws:

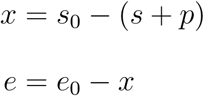

This model was used to fit the elastase kinetic data (Figure 2), as described in the fit methods section. Table 3 reports the results of our best-fit, which seem consistent with the results of the fit with the full three-step model, reported in the Supporting Information (S.I.).

**Table 3:**
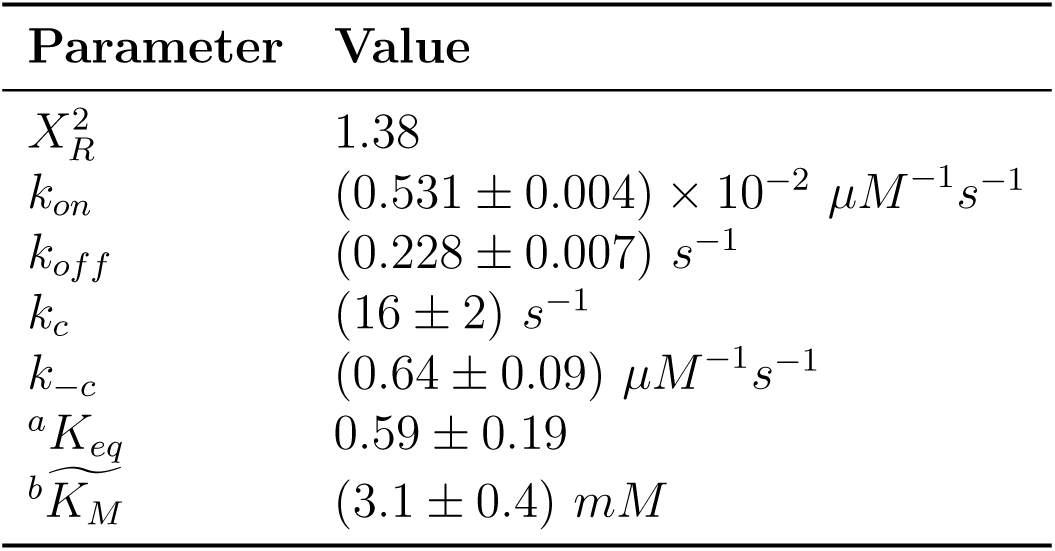
Best-fit values for the Michaelis Menten model. *^a^ K_eq_* = *k_on_k_c_*/*k_off_ k_−c_*; *^b^*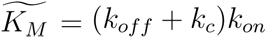.

| **Parameter** | **Value** |
|---|---|
| *X*<sub>*R*</sub><sup>2</sup> | 1.38 |
| *k*<sub>*on*</sub> | (0.531 ± 0.004) × 10<sup>−2</sup> *µM*<sup>−1</sup>*s*<sup>−1</sup> |
| *k*<sub>*off*</sub> | (0.228 ± 0.007) *s*<sup>−1</sup> |
| *k*<sub>*c*</sub> | (16 ± 2) *s*<sup>−1</sup> |
| *k*<sub>−*c*</sub> | (0.64 ± 0.09) *µM*<sup>−1</sup>*s*<sup>−1</sup> |
| <sup>*a*</sup>*K*<sub>*eq*</sub> | 0.59 ± 0.19 |
| $^{b}\widetilde{K_M}$ | (3.1 ± 0.4) *mM* |

First of all, it must be noted that the absolute values resulting from the fits are affected by the imperfect agreement between the two experiments performed for the 750 µM time series. The value of *k_c_* = 16 *s^−^*^1^ is in good agreement with most of the values reported in Table 2 for *k_cat_*. The value for *k_off_* is very close to one reported by Nunes et al. for the dissociation of the intermediate complex formed by HNE with a FRET-peptide [37]. We are not aware of previous estimations of *k_−c_* and of the equilibrium constant *K_eq_* = *k_on_k_c_*/*k_off_ k_−c_*. The latter suggests that at equilibrium, 36 % of the initial substrate is converted into products. The estimate of 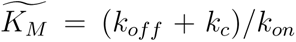 appears inconsistent with the values reported in Table 2 for *K_m_*. This is likely because the inclusion of the reversible step prevents the direct identification of 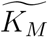 with *K_m_*. The best-fit value of the association rate *k_on_* turns out to be (0.531 *±* 0.004) *×* 10*^−^*^2^ µ*M ^−^*^1^*s^−^*^1^, roughly 6 orders of magnitude lower than the typical diffusion-limited rate of enzymatic reactions, which is around 10^4^ µM *^−^*^1^*s^−^*^1^ [42], and one order of magnitude lower than the best-fit value of the product recombination constant *k_−c_*. In the S.I. we show that a rate-limiting conformational step, of the kind previously suggested by Stein and collaborators [30, 31, 43], can justify this apparently harsh slow-down of the association step in the coarse-grained scheme.

### The Michaelis Menten scheme versus alternative models

An irreversible MM model cannot reproduce the long-time behaviour of our data by construction. In fact, for long time series of product formation approaching equilibrium, it is necessary to adopt a thermodynamically consistent model featuring microscopic reversibility. However, a simple reversible MM model (11) with an association constant *k_on_* close to the diffusion-limited value is equally unable to correctly reproduce the measured time-courses.

We found that the use of a reversible three-step mechanism of serine proteases slightly improved the results, but did not allow to capture the kinetic behaviour at all substrate concentrations simultaneously. As already anticipated, this does not come as a surprise, because our data stretch to the long-term regime of the catalysis, hardly distinguishing between the two catalytic constants of serine proteases (see the S.I. for further details).

On the other hand, the data can be fitted with a single set of the rate constants simultaneously for all substrate concentrations by models that include conformational steps for the intermediate complexes. One of the simplest hypotheses is that a single intermediate conformational step is required to generate a productive complex. This is reminiscent of the hypothesis of a rate-limiting conformational step preceding acylation and deacylation, formulated by Stein and colleagues to account for the peculiarity of the experimental observation of the hydrolysis of the M-pNA peptide by HNE [30, 31, 43]. In the S.I. it is shown how a simplified version of this model, where enzyme acylation and deacylation are accounted for through a single catalytic step, leads to effective rate-limiting *k_on_* and *k_off_*, with a fast association step k_1_, closer to the diffusion-limited rate.

## 6. Effect of crowding on the kinetics of HNE

The simplest assumption is that crowding does not alter the kinetic scheme that describes the catalytic conversion, but simply renormalizes the rate constants to concentration-dependent values. Accordingly, we assume that the one-step MM reversible model still provides a faithful interpretative paradigm. This cannot be given for granted in general, as crowding might lead to the breakdown of some of the hypotheses underlying the procedure outlined in section 5, e.g. the homogeneity of the solution.

All the experimental time courses and the relative MM fits are illustrated in Fig. 4. The corresponding best-fit values of the fitting parameters are plotted in Fig. 5 versus crowding concentration. It should be emphasized that DEX10 time series are additionally corrected by dextran absorption, as required by our calibration procedure (section 3.1). Interestingly, the kinetic data collected in the presence of DEX450, exhibit a complex trend at short times, notably at high concentration. This cannot be accounted for within a simple MM scheme. In fact, the reaction velocity is seen to increase with time and go through a maximum (see Fig. 4i, 4l, 6 and S6). In such cases, we estimate the time *t_M_* corresponding to the maximum velocity, *V_M_*, and then fit the data as if the reaction had started with constant maximum velocity at time t_0_ = *P*(*t_M_*)/*V_M_ − t_M_*. Only the points at *t* > *t_M_* are then included in the fit. This procedure allows us to consistently extract information on the kinetics of HNE in the presence of DEX450, at least at low and moderate concentrations.

**Figure 5:**
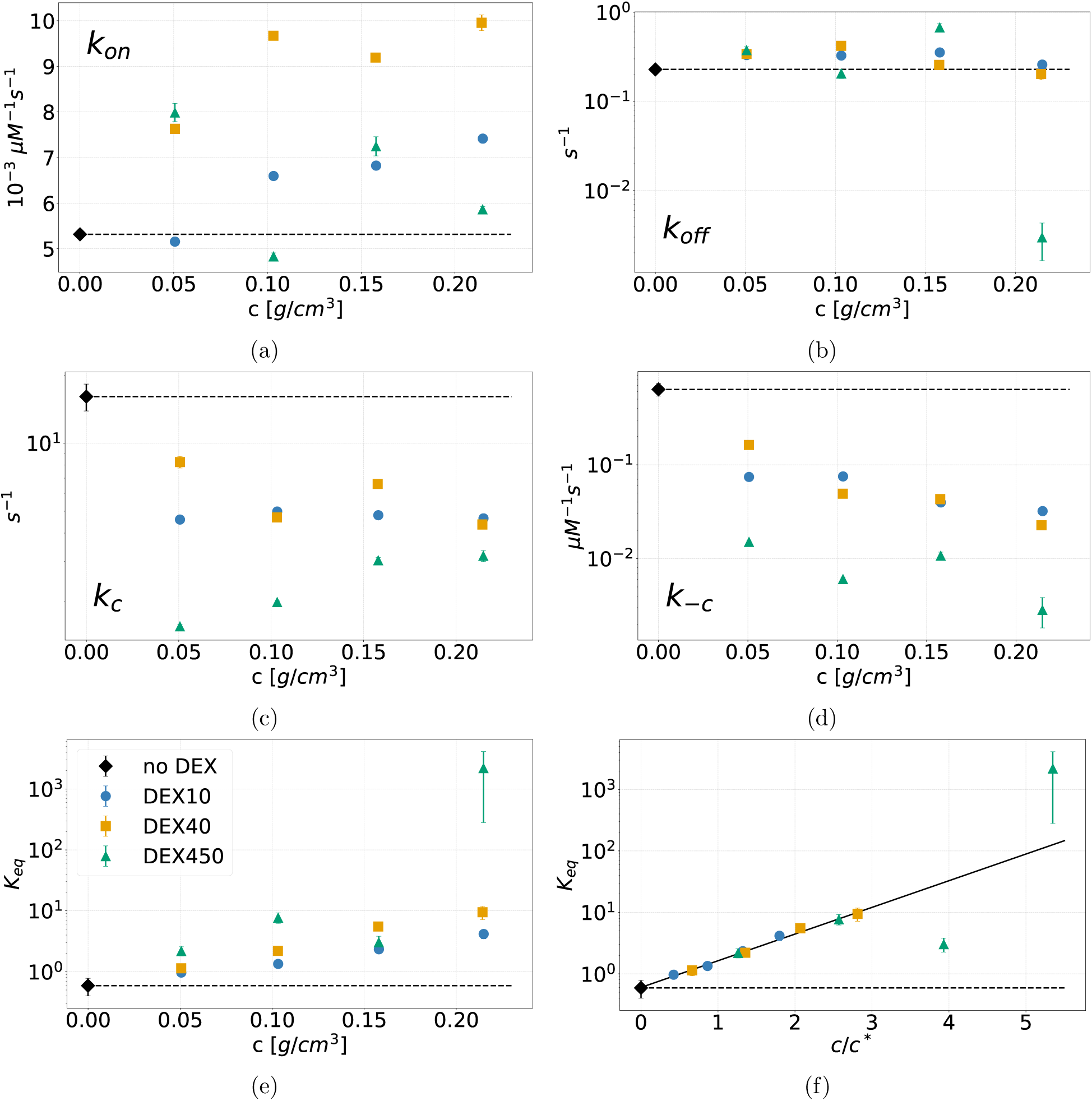
Best-fit values of the reversible MM kinetic constants vs crowding concentration. Panel (f) shows the equilibrium constant as a function of the scaled variable *c/c^∗^*.

**Figure 6:**
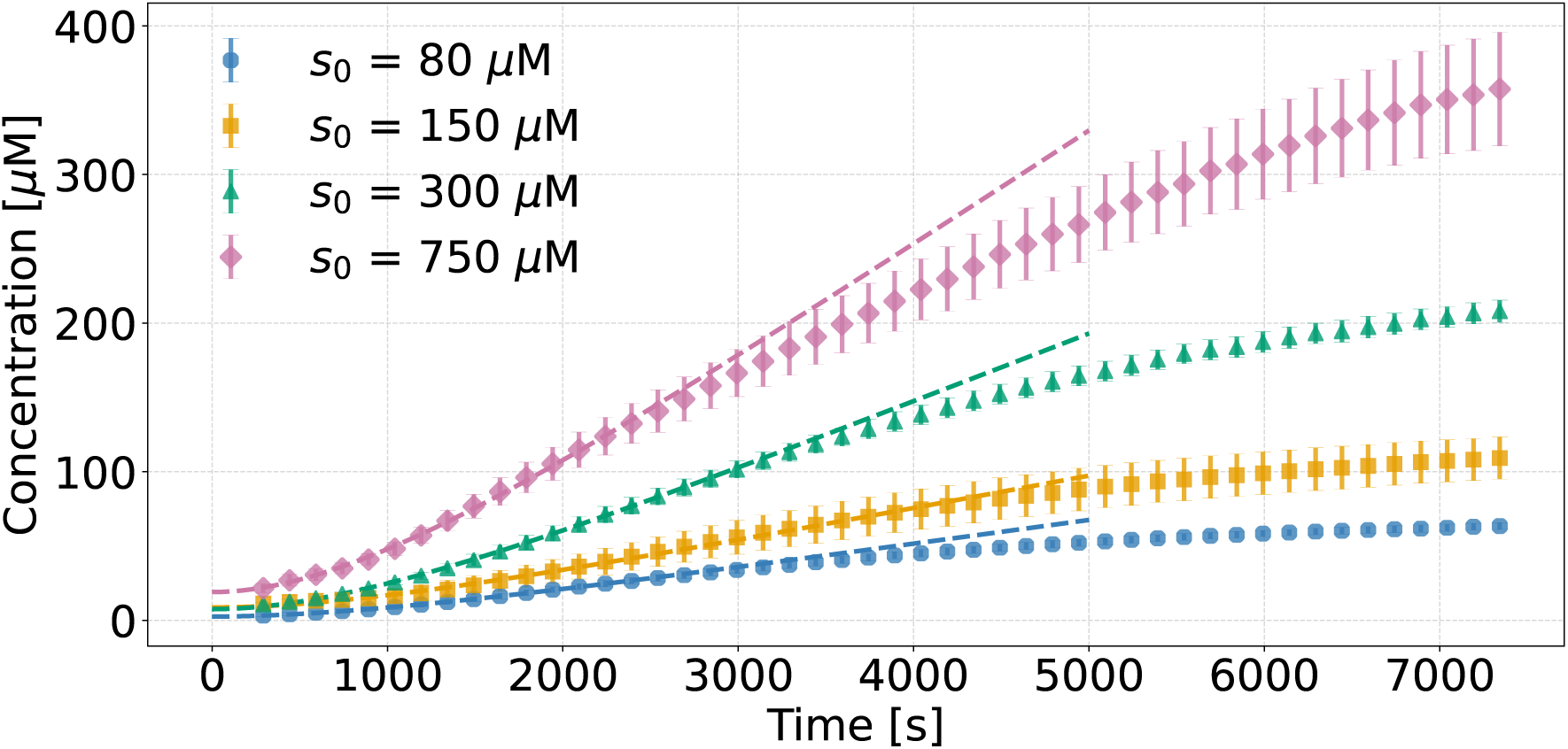
Product generation in the presence of DEX450 at 20 % weight fraction. The data shown and the error bars represent averages and standard errors computed over 4 replicates. The dashed lines are simultaneous fits to Eq.(18) up to T_max_ = 2500 s, with a common relaxation timescale τ = 951 *±* 55 s.

Our fitting procedure reveals that *k_c_* and *k_−c_*, the rate constants associated with enzyme acylation, deacylation and recombination of the product with the free enzyme, decrease with crowding concentration (see Fig. 5c and 5d). The complex dissociation rate, *k_off_* takes values in the range 0.20 *−* 0.45 *s^−^*^1^. The association rate, *k_on_*, increases non linearly with concentration for DEX10 and 40, suggesting that this constant is approaching a limiting value. This behaviour is reminiscent of a prediction for the constant associated with the formation of the first intermediate complex, *k*_1_ [44], which in our case is proportional to *k_on_* (see the S.I.).

Interestingly, the variation of the equilibrium constant *K_eq_* = *k_on_k_c_*/ (*k_off_ k_−c_*) leads to interesting insight. Figs. 5e and 5f show that *K_eq_* increases with concentration somewhat differently, depending on the size of the crowder. The increase of *K_eq_* marks an increasing shift of the reaction equilibrium towards the product, up to 10 times more at high crowding concentration than in the dilute regime. Motivated by an analogy of such trends with well-known scaling properties of polymer physics [15, 16] found by other authors in similar dextran systems [21] and by us in the same samples analyzed here [24], we posit that *K_eq_* is described by a scaling form of the kind

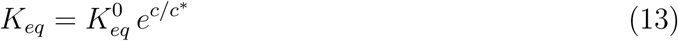

where *c^∗^* is the so-called overlap concentration [17], i.e. the typical relevant scaling concentration for viscosity, self-diffusion and osmotic pressure in polymer physics [15, 17, 21]. The zero-crowding value 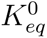 is fixed by the best estimate obtained in the absence of crowders, 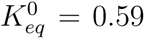. All (four) data points, in the presence of either DEX10 or DEX40, are well reproduced by Eq. (13) with the following estimate for the overlap concentration *c^∗^*: 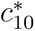 = 0.119 ± 0.004 g/cm^3^ and 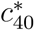 = 0.076 ± 0.002 g/cm^3^. For DEX450, scaling appears to restrict reasonably to the two lower concentrations, with 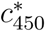 = 0.0402 ± 0.0004 g/cm^3^. As shown in Figure 5f, the exponential function (13) captures to a very good extent the trend of *K_eq_* with concentration. The quantity *c^∗^* is expected to scale with the molecular weight M as *c^∗^ ∼ M ^−a^*, where a = 3ν *−* 1 and ν is the Flory exponent [17]. Therefore, from 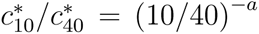, we get a *≈* 0.32, which corresponds to a Flory exponent ν = (1 + a) /3 = 0.44. Analogously, one can estimate *a* and *ν* through 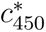 and 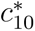 or 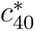.

The results of these calculation are summarized in Table 4. The Flory exponent ν *≃* 0.44 is in good agreement with the experimental values for dextran available in the literature; for instance, with estimates from scaling properties of the second virial coefficient [22, 45], of the Stokes-Einstein radius [46], the mean radius of gyration and the translational diffusion [22]. Interestingly, this value is also very close to the Flory theory for randomly branched polymers in θ*−*solvent [47]. In addition, simulations of branched quenched polymers often report the estimations of ν lower than 0.5 [48, 49].

**Table 4:** Estimates of the exponents *a* and *ν* = (1 + *a*) /3 obtained by comparing pairwise the overlap concentrations from the exponential fits of *K_eq_* (see Fig. 5f). For DEX450, we used 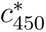 = 0.0402 *±* 0.0004 g/cm^3^.

| Sizes compared [kDa] | *a* | *ν* |
|---|---|---|
| 10 − 40 | 0.32 | 0.44 |
| 10 − 450 | 0.29 | 0.43 |
| 40 − 450 | 0.26 | 0.42 |

From a statistical mechanics perspective, the equilibrium constant *K_eq_* can be divided into an ideal and a non-ideal contribution [6, 44]

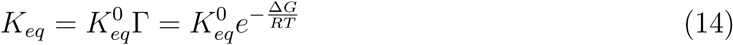

where Γ is a factor that accounts for non *−* ideal contributions to the Gibbs free energy of the reaction Δ*G* = *G(P)* − *G(S)*. A comparison with Eq. (13) suggests that Δ*G* is a linear function of concentration, which scales with the overlap concentration *c^∗^* as

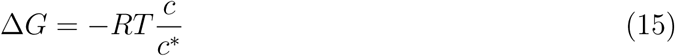

The result of the exponential fits can be therefore interpreted in the following way. Macro-molecular crowding by dextran leads to a decrease of the non-ideal Gibbs free energy of the enzymatic reaction, which is linear in dextran mass concentration. This amounts to a shift towards the product in the reaction equilibrium. This conclusion is valid up to concentrations of the order 0.2 g/cm^3^, for DEX10 and DEX40, and roughly 0.10 g/cm^3^ for DEX450. Fig. 5f suggests that the exponential scaling holds, independently from dextran size, up to *c/c^∗^ ≃* 3. This range of validity is not different from the range of universal behaviour found by Burchard and coworkers for osmotic modulus and viscosity [21]. Concerning the peculiar behaviour of DEX450, one may note that HNE has a molecular mass around 25 kDa, that is, of the same order as polysaccharides such as DEX10 and DEX40, while DEX450 is significantly larger.

It is interesting to observe that, in the literature on macromolecular crowding, crowders are often approximated as hard spheres, ellipsoids or other simple geometrical forms [44, 50]. Typically, this involves scaled-particle theory [51] or the concept of available volume [6], even though the volume associated with macromolecules in solution is not easy to define [6] and the approximation of the crowder as a simple geometry is likely inaccurate for a flexible polymer [11, 50]. In our reasoning, we do not make any *a priori*assumption on the molecular shape of crowders. This allows us to identify the scaling parameter *c^∗^* as the overlap concentration, a quantity characteristic of polymer physics that incorporates implicitly information on the topology and the interactions with the solvent, through the Flory exponent ν. These considerations suggest that using concepts from polymer physics when interpreting crowding results may lead to relevant insight, notably for polymer crowders, as recently observed also by other authors [52, 53]. For instance, the quantity *c/c^∗^* can be interpreted as the packing fraction of spheres with radius *R_g_*, the gyration radius of polymers in dilute conditions. However, for flexible polymer systems the ratio *c/c^∗^* cannot be directly traced back to the volume available to an infinitesimal tracer. In our case, this is mainly due to the soft nature of dextrans, which are flexible polysaccharides due to the prevalence of α(1 *→* 6) glycosidic bonds [21, 54]. Therefore, our results reinforce the importance of accounting for the softness of crowders beyond the hard-sphere approximation as also evidenced in other works [55, 56].

The dextran-dependent shift in the inferred equilibrium constant raises a central mechanistic question: does crowding alter the thermodynamic balance between substrate and products, or does it modify enzyme dynamics over the experimental observation window? Macromolecular crowding can induce conformational changes that influence enzymatic activity [2] and equilibria involving macromolecular species [6, 7]. Because the reaction did not reach an experimentally resolved equilibrium, we cannot exclude the possibility that the fitted K_eq_ captures a transient regime involving elastase. In this scenario, crowding-induced stabilization of a more active enzyme conformation could contribute to the apparent shift.

The substantial decrease in reaction velocity towards the end of the experiments (Figures S5 and S6) is consistent with an approach to equilibrium, although it does not uniquely establish this interpretation. If the inferred K_eq_ represents the reaction equilibrium, its dependence on dextran concentration must reflect changes in the relative activities of the free substrate and products. Enzyme conformations and catalytic intermediates can affect the rate of approach to equilibrium, but their contributions cancel from the thermodynamic balance of the overall catalytic reaction. The resulting shift would therefore implicate differential interactions of dextran with substrate and products, either directly or through changes in their solvation and ionic environment.

Depletion interactions have been invoked to explain crowding effects in polymer solutions [57, 58], but their applicability to the small substrate and products studied here requires further scrutiny. An additional possibility follows from the change in charge accompanying hydrolysis. Previous studies report that dextran and other common crowding agents can associate with sodium ions, potentially modifying the local ionic environment [59]. Whereas MeoSuc-AAPV-AMC is neutral, cleavage exposes a C-terminal carboxylate on the peptide product, which is expected to be negatively charged at physiological pH. Differential electrostatic and solvation effects could therefore contribute to the observed shift. Testing the dependence of the scaling on ionic strength and product charge would help distinguish this mechanism from other contributions to the crowding response.

Finally, our results suggest that, in systems crowded with polymers, experiments with species of different sizes might be understood in terms of the scaled variable *c/c^∗^*, as highlighted also in our recent study on physical properties of dextran solutions [24]. In this context, performing the experiments with polymers of different topology, we expect *c^∗^* to scale with size according to the appropriate Flory exponents. If the effect of crowding on an enzymatic reaction is dominated by the interaction between the solvent and the monomer of the polymeric crowder, no dependence from the size is expected instead. The importance of accounting for the structure of the polymer also stems from the observation that, in the case of hard spheres, ν = 1/3 and consequently *c^∗^ ∼ M* ^1^*^−^*^3^*^ν^* do not depend on the size.

### HNE kinetics in the presence of DEX450

Product generation kinetics in the presence of DEX450 displays a peculiar nonlinear behaviour at short times. This is illustrated in Fig. 6. Moreover, our analysis of the asymptotic regime shows that the rate constants at high concentrations deviate from the trends observed for DEX10 and DEX40 (see Fig. 5). This suggests that these solutions behave in a significantly different manner at high concentration with respect to the smaller polymers [45]. There certainly are some specific experimental aspects to be considered as the simplest explanation first. In our protocol, the first minutes of the reaction after substrate addition to the solutions cannot be recorded through the plate reader, because of the mixing. One might surmise that, due to the high viscosity of DEX450 solutions at high concentration, the 3 minutes-long mixing might not be enough to ensure homogeneity. As the substrate is the last reactant added to the solution, there might be isolated droplets left after addition that require more time to dissolve in the solution. If the cause of the transient effect were not purely experimental, more complex reaction mechanisms should be surmised to explain this observation. However, there is no indication of alternative or additional kinetic steps induced by the interaction with DEX450. Hence, it is preferable to look for an explanation that does not require unjustifiably complex biochemical networks.

In the slow-mixing hypothesis, the slow dynamics would reflect an additional rate-limiting phenomenon to be traced back to the properties of DEX450 solutions. These are likely heterogeneous at the microscopic scale, which might hamper substantially enzyme-substrate diffusive encounters. A simple way to account for a slow mixing kinetics that transiently affects the formation of enzyme-substrate complexes is to divide the enzyme pool in two populations at a coarse-grained level. Let us imagine a catalytically poorly accessible state *E_p_* and a catalytically accessible state *E_a_*, with 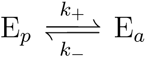. This notation should not be interpreted literally as a specific microscopic conformational transition. In a crowded dextran solution, *E_p_* and *E_a_* may represent effective ensembles that differ in spatial partitioning and/or local substrate environment and accessibility. Let *E_T_* = *E_p_* + *E_a_* denote the total concentration of enzyme, constant during the fitted time window. The data indicate the maximal-lag limit, in which the initially accessible catalytic activity is small compared with its stationary value. In that limit, one has

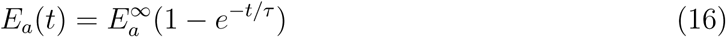

with τ = 1/(*k*_+_ + *k_−_*) and 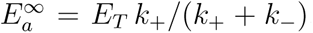. It is reasonable to assume that binding, unbinding, and catalytic turnover equilibrate sufficiently rapidly relative to the slow accessibility relaxation. Under this separation of time scales, at any instant, the productgeneration rate is proportional to the currently accessible catalytic capacity. Therefore, at initial substrate concentration *S*_0_, we may write

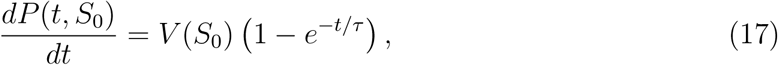

where *V* (*S*_0_) is the asymptotic velocity that would be reached after the slow relaxation has completed, within the regime in which substrate depletion remains negligible. These should be expected to be linear with S_0_ at low substrate concentration and saturate to a maximal velocity at higher concentrations, according to the MM prescription for initial velocities. Integrating Eq. (17) from *t* = 0 to *t* = *∞* and introducing curve-specific additive nuisance parameters *P*_0_(*S*_0_), we get

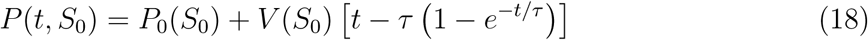

The parameters *P*_0_(*S*_0_) are *not* to be interpreted as literal concentrations of product present at the physical start of the reaction. The first genuine measurements occur at approximately 292 s, so these are backward-extrapolated intercepts. Their values may reflect the heterogeneous mixing process, without implying a non-zero non-monotonic product concentration at the experimental start.

As Fig. 6 demonstrates, the data can be fitted simultaneously up to *T*_max_ = 2500 s through Eq. (18) with a single slow relaxation time scale, τ = 951 *±* 55 s, for DEX450 at 20% weight fraction. Importantly, the four fitted values of V (S_0_) appear to follow a MM trend, as expected (see the S.I.). This analysis can be repeated for all the considered weight fractions of DEX450 with the same results. Interestingly, τ is found to increase monotonically with DEX450 weight fraction, which is consistent with our hypothesis of crowding-induced slow relaxation phenomenon (see the S.I.).

## 7. Conclusions

In this work, we studied the degradation of a short peptide by elastase with fluorometric assays in different solutions crowded with dextran polymers. An innovative calibration method introduced recently by us [32] was applied to study the reaction at high concentrations of substrates and crowders.

For each dextran size and concentration, the enzymatic kinetic data were recorded and fitted simultaneously for four different initial substrate concentrations. A good quality of the fit was found using a reversible MM mechanism, with one single intermediate complex. It was not necessary to consider separately a sequential release of reaction products, as in the archetypal model of serine proteases [26], likely because the reaction was observed in a time range much longer than the characteristic times typically associated with the release of the products (see Table 2). However, we found a low value of the association rate constant, *k_on_ ∼* 0.5 *×* 10*^−^*^2^ µ*M ^−^*^1^*s^−^*^1^, incompatible with the diffusion-limited rate. We interpreted this result as possible evidence of a rate-limiting conformational step preceding the formation of the catalytically competent complex. A similar hypothesis had been introduced by Stein and coworkers in another context [30, 31, 43].

In the presence of dextran, for all the considered sizes and concentrations, the total product generated by HNE over the two hours of measurement was found to increase with crowding. With the exception of the data involving the higher concentrations of DEX450, the kinetic activity could be interpreted with the same model used in absence of crowders. It was possible to estimate the equilibrium constant of the reaction, *K_eq_*, which was found to increase with dextran concentration in a size-dependent way. Each curve could be fitted to a good extent with an exponential, introducing a size-dependent scaling concentration, *c^∗^*, which led the equilibrium constants to collapse on a universal curve.

We suggest to identify *c^∗^* with dextran overlap concentration. Indeed, the values estimated from the fits scale with dextran molecular weight M as M ^1^*^−^*^3^*^ν^* with Flory exponent ν *≃* 0.44. This value is recurrent in studies of dextran solutions [21] and is typically interpreted as a sign of branching [22, 45]. We also found the same value observing the scaling properties of self-diffusion and viscosity [24].

While a size dependence in the crowding effect on an enzymatic reaction may be interpreted as a blueprint of inter-molecular interactions, our results suggest that the *topology* of crowders may be used as a tunable parameter besides their size. In fact, for an enzymatic reaction whose equilibrium properties depend on the size of the polymeric crowder, as in our case, we expect the scaling properties to be determined by the polymer topology, through the size-dependence of *c^∗^*.

Overall, our study clearly demonstrates the importance and predictive power of interpreting the effects of crowding realized with polymers on purely thermodynamic grounds, i.e. without assimilating the crowders to hard spheres or ellipsoids as often done in the field of macromolecular crowding.

## Acknowledgements

Enlightening discussions with Anand Yethiraj are gladly acknowledged by all authors. G.M. thanks collaborators and PhD students at Centro di Studio Dinamiche Complesse (University of Florence) and Centre de Biophysique Moléculaire for stimulating conversations. G.M. is also grateful to Robert Hołyst for insightful discussions on the role of charges and ionic strength in polymer solutions. This work was supported by the French Agence Nationale de la Recherche (ANR), through the project X-CROWD, contract ANR-21-CE44-0020-01.

## Author contributions

F.P. and J. H. designed the study and supervised the theoretical and experimental activities, respectively. J.C.V. and G.M. contributed equally to this work as co-first authors. J.C.V. conducted all experimental work. G.M. developed the computational tools, analysed the data and contributed substantially to their interpretation. All authors contributed to discussing the results and writing the manuscript.

## Footnotes

∗ In order to calculate the molar concentration of DEX10 the value of molecular weight used was taken from the manufacturer and for DEX10 it was 9126 g/mol.

