## Supporting Information for "Polymer overlap and the organization of enzyme activity in crowded solutions"

#### Contents

**1 Fluorescence calibration** **S-3**

1.1 Fluorophore and substrate . . . . . . . . . . . . . . . . . . . . . . . . . . . . . S-3

1.2 Fluorophore in the presence of Dextran . . . . . . . . . . . . . . . . . . . . . . S-3

**2 Reaction velocity** **S-7**

**3 Kinetic models and fits in the absence of crowders** **S-9**

3.1 Three-step model . . . . . . . . . . . . . . . . . . . . . . . . . . . . . . . . . . S-9

3.1.1 Quasi steady-state approximation . . . . . . . . . . . . . . . . . . . . . S-9

3.2 Two products kinetic model . . . . . . . . . . . . . . . . . . . . . . . . . . . . S-13

**4 Slow-homogeneisation model of kinetics in the presence of DEX450 S-15**

### 1 Fluorescence calibration

#### 1.1 Fluorophore and substrate

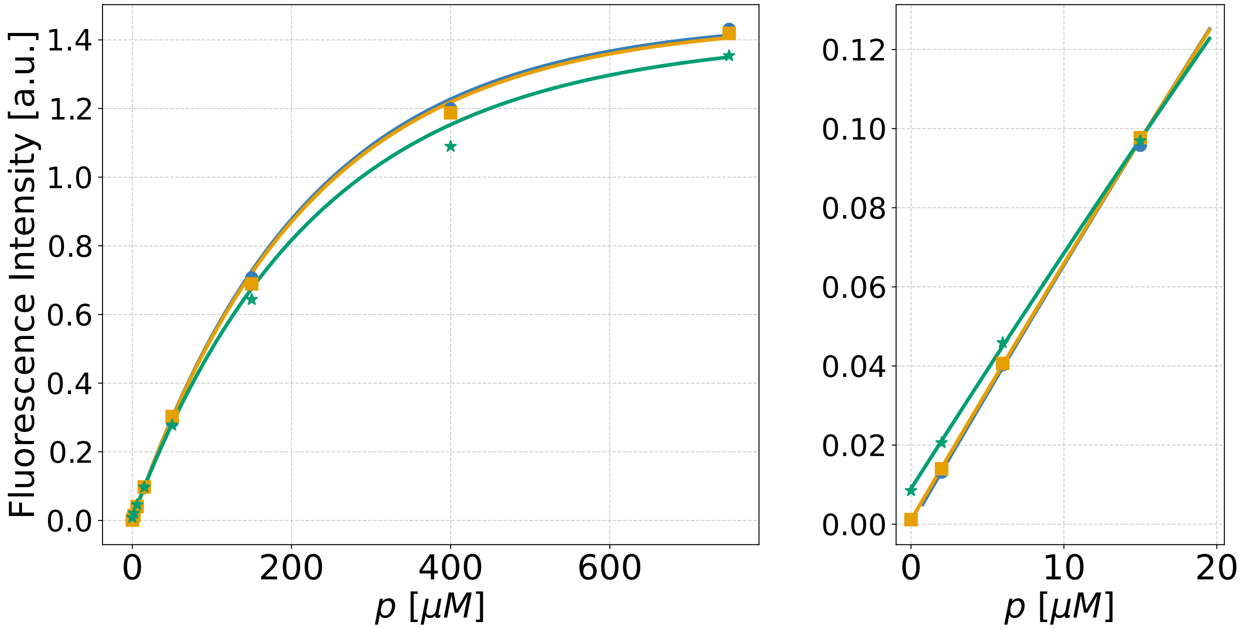


Figure S1: Calibration curves for the fluorescence intensity (FI) *vs.* the product concentration (AMC fluorophore) recorded at different values of the substrate (AMC-peptide) concentration $s$. Symbols and colors mark the measured fluorescence: $s = 0$ , blue circles; $s = 80$ $\mu$M, orange squares; $s = 750$ $\mu$M, green stars. Solid lines correspond to the fit performed via Eq. (4) in the main text, in the absence of crowders. The right panel shows a close-up of the measured data in the diluted regime, along with the linear approximation of the theory, $F = \alpha(p + rs)$. The small effect of the residual emission at high substrate concentration (green stars) shows up clearly.

#### 1.2 Fluorophore in the presence of Dextran

Calibration of AMC in the presence of dextran. Fits are performed according to relation:

$$F = \frac{\alpha p}{\zeta p + \zeta_D c_D} \left(1 - e^{-(\zeta p + \zeta_D c_D)}\right) \tag{1}$$

Here $\zeta_D$ and $c_D$ refer to the absorption coefficient and concentration of dextran. Differently from the case of DEX10, reported in Figure SI.S2, extinction could not be meaningfully resolved from zero for DEX40 and DEX450, as shown in Figures S3 and S4, and was neglected.

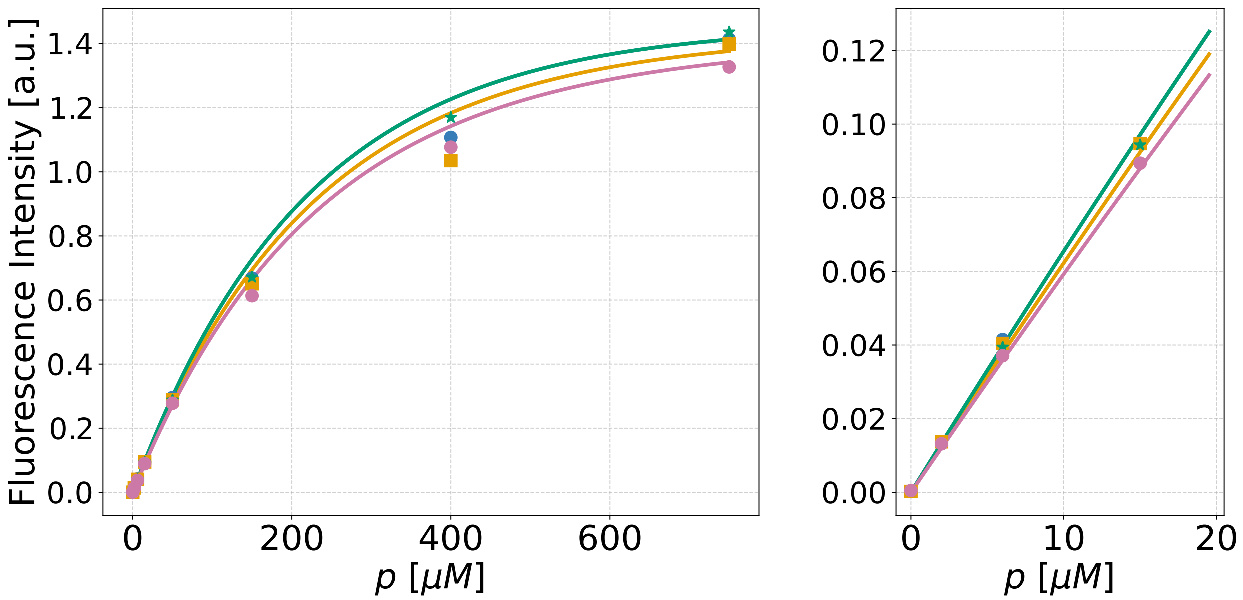


Figure S2: Calibration curves for the fluorescence intensity (FI) *vs.* the product concentration (AMC) for different values of DEX10 concentration $c_{10}$. Symbols and colors mark the measured fluorescence: $c_{10} = 0$ , blue circles and orange squares; $c_{10} = 1.12 \times 10^4$ $\mu$M (0.10 w/w), green stars; $c_{10} = 2.24 \times 10^4$ $\mu$M (0.20 w/w), pink circles. Solid lines correspond to the fit performed via Eq. (4) in the main text, with $\alpha$ and $\zeta$ fixed to the value in the absence of crowders. The best estimate of the parameter $\zeta_{10}$ obtained through this fit is reported in Table 1 in the main text. The right panel shows a close-up of the measured data in the diluted regime, along with the linear approximation of the theory, $F = \alpha p$. No residual emission at high dextran concentration is observed.

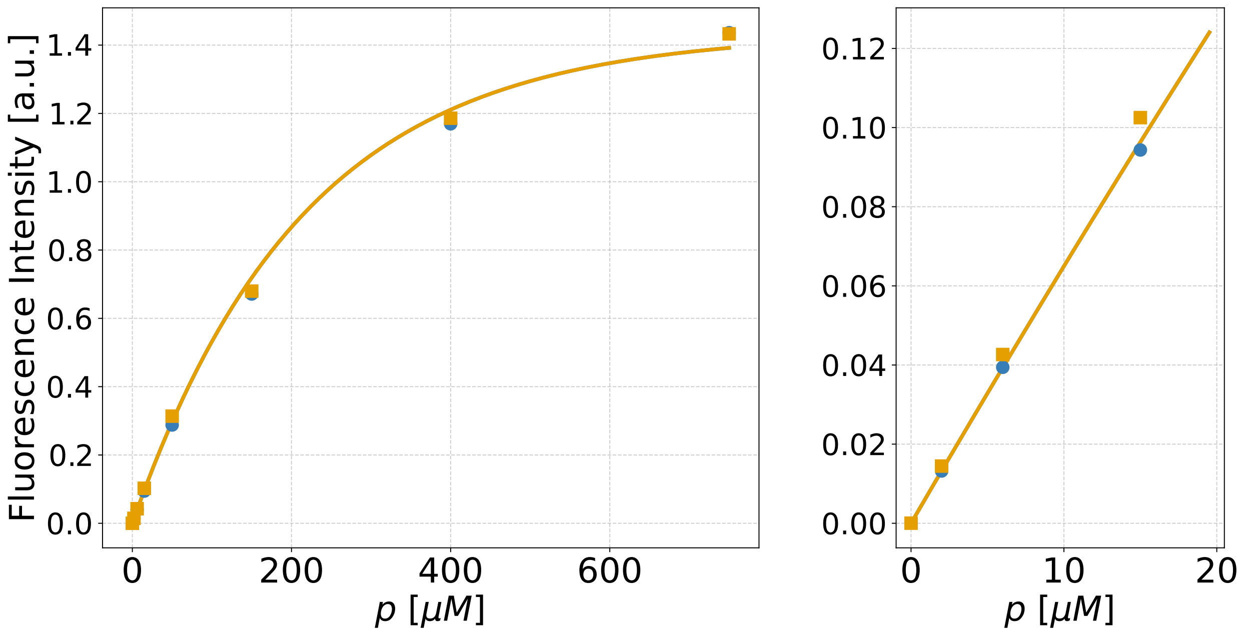


Figure S3: Calibration for the product (AMC) for different values of DEX40 concentration $c_{40}$. Symbols and colors mark the measured fluorescence: $c_{40} = 0$ , blue circles; $c_{40} = 0.56 \times 10^4$ $\mu$M (0.30 w/w), orange squares. $c_{10} = 2.24 \times 10^4$ $\mu$M (0.20 w/w), pink circles. Solid lines correspond to the fit performed via Eq. (1), with $\alpha = (0.665 \pm 0.010)$ $\mu M^{-1}$, $\zeta = (4.6 \pm 0.2) \times 10^{-3}$ and $\zeta_{40} = (0.0 \pm 0.9) \times 10^{-5}$.

The conversion of the fluorescence intensity into concentration units requires to solve numerically the equation

$$F_n - F(p_n; \vec{\alpha}) = 0 \tag{2}$$

for each point in the fluorescence data $F_n$. The calibration relation $F(p_n; s_0, \vec{\alpha})$, eq. (10) in the paper, is defined through the vector of parameters $\vec{\alpha} = (\alpha, r, \zeta, \gamma, \zeta_{10})$. $s_0$ is also a parameter in the inversion, to which we associate an error of $e_0/2$, the width of the interval in eq. (9) in the paper. $F(p_n)$ is a monotonic function of $p_n$ and therefore its inverse is well defined

$$p_n = F^{-1}\left(F_n; s_0, \vec{\alpha}\right) \tag{3}$$

Although $F^{-1}$ cannot be expressed in an explicit manner, the function is well defined numerically and with standard error propagation techniques, we express the uncertainties on

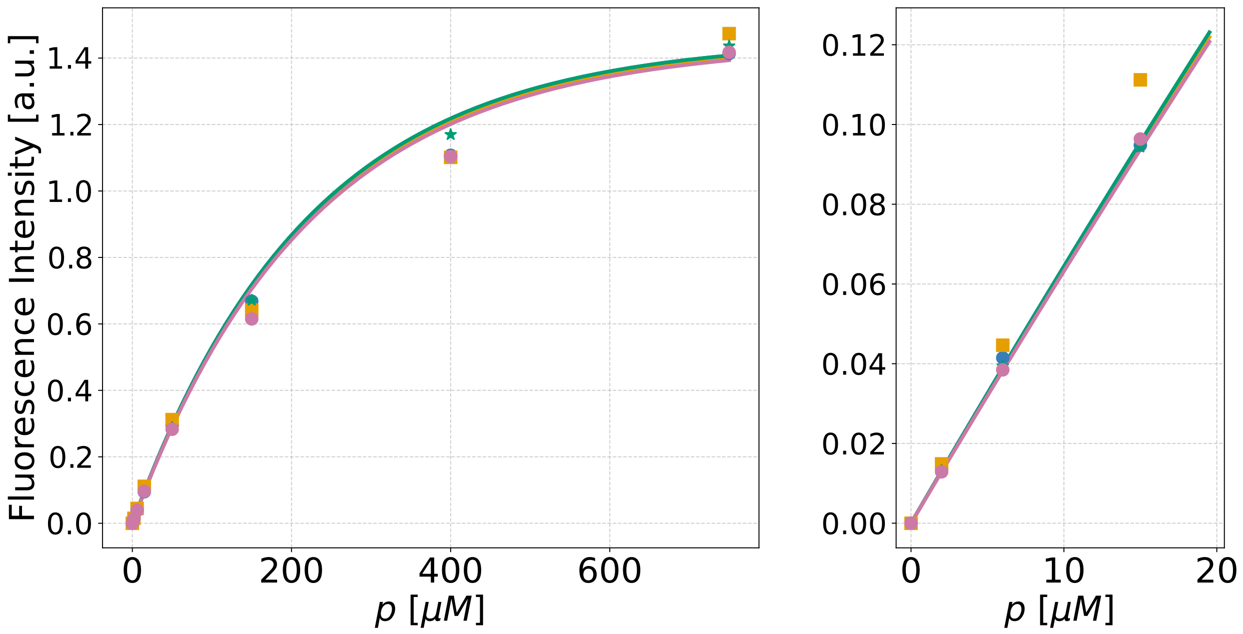


Figure S4: Calibration for the product (AMC) for different values of DEX450 concentration $c_{450}$. Symbols and colors mark the measured fluorescence: $c_{450} = 0$ , blue circles and orange squares; $c_{450} = 2.16 \times 10^2$ $\mu$M (0.10 w/w), green stars; $c_{450} = 4.32 \times 10^2$ $\mu$M (0.20 w/w), pink circles. Solid lines correspond to the fit performed via Eq. (1),with $\alpha = (0.659 \pm 0.005)$ $\mu M^{-1}$, $\zeta = (4.52 \pm 0.09) \times 10^{-3}$ and $\zeta_{450} = (0.9 \pm 1.1) \times 10^{-4}$.

$p_n$ as

$$\sigma_n = \frac{1}{|\frac{\partial F}{\partial p_n}|} \sqrt{\Delta F_n{}^2 + \left(\frac{e_0}{2}\frac{\partial F}{\partial s_0}\right)^2 + \sum_{\alpha_i,\alpha_j} cov(\alpha_i,\alpha_j)\frac{\partial F}{\partial \alpha_i}\frac{\partial F}{\partial \alpha_j}} \tag{4}$$

where $\Delta F_n$ are the standard errors on $F_n$, estimated over the 4 replicates. The covariance matrix of the calibration relation is directly obtained as the result of the fit over the binary mixtures:

$$\mathrm{Cov}\,(\alpha_\mathrm{i}, \alpha_\mathrm{j}) = \begin{pmatrix} 1.63 \times 10^{-5} & -3.63 \times 10^{-8} & 1.15 \times 10^{-7} & 7.50 \times 10^{-6} & 0 \\ -3.63 \times 10^{-8} & 5.28 \times 10^{-9} & -1.07 \times 10^{-9} & 2.73 \times 10^{-7} & 0 \\ 1.15 \times 10^{-7} & -1.07 \times 10^{-9} & 2.40 \times 10^{-9} & -1.80 \times 10^{-7} & 0 \\ 7.50 \times 10^{-6} & 2.73 \times 10^{-7} & -1.80 \times 10^{-7} & 8.08 \times 10^{-5} & 0 \\ 0 & 0 & 0 & 0 & 8.30 \times 10^{-11} \end{pmatrix}$$

### 2 Reaction velocity

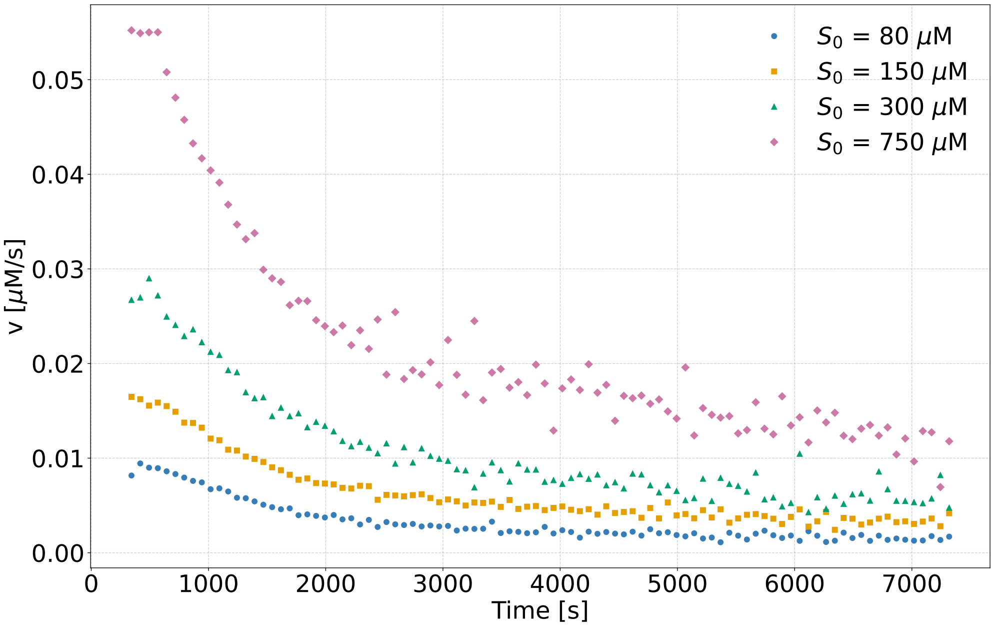


Figure S5: Reaction velocity as a function of time in the absence of crowders, estimated as simple finite difference of the product time series.

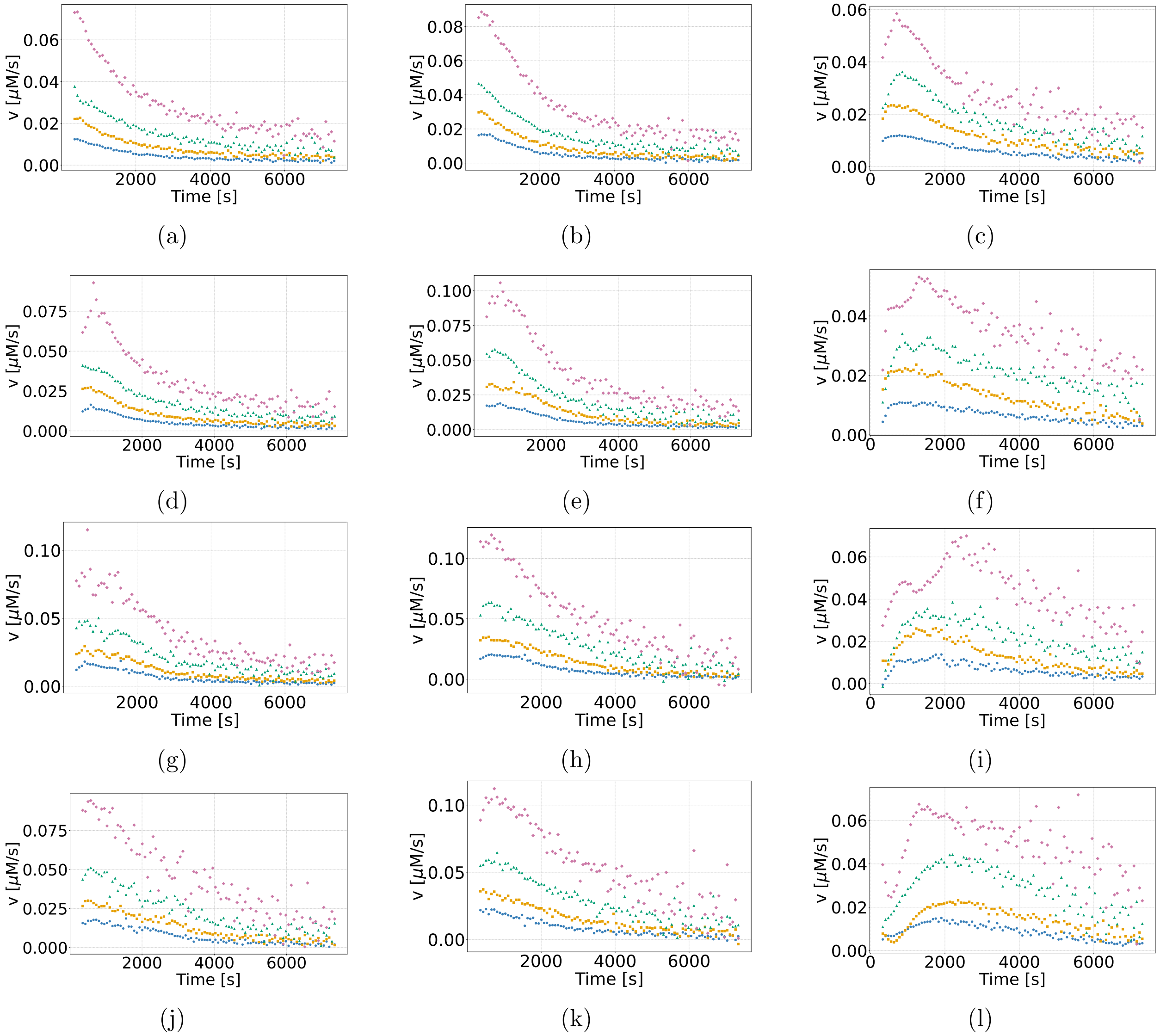


Figure S6: Reaction velocity in the presence of dextran at different weight fractions $w$. In columns, from left to right, DEX10, DEX40 and DEX450 can be observed. In rows, dextran weight fraction is fixed, from top to bottom, dextran weight fractions is: 0.05, 0.10, 0.15 and 0.20. DEX450 shows a transient lag time before reaching maximum velocity.

### 3 Kinetic models and fits in the absence of crowders

#### 3.1 Three-step model

Let us consider the following model

$$\mathrm{S} + \mathrm{E} \underset{\mathrm{k}_{-1}}{\overset{\mathrm{k}_1}{\rightleftharpoons}} \mathrm{X}_a \underset{\mathrm{k}_{\mathrm{ba}}}{\overset{\mathrm{k}_{\mathrm{ab}}}{\rightleftharpoons}} \mathrm{X}_b \underset{\mathrm{k}_{-c}}{\overset{\mathrm{k}_c}{\rightleftharpoons}} \mathrm{E} + \mathrm{P} \tag{5}$$

This scheme gives rise to the following system of ODEs for the concentrations

$$\begin{aligned} \frac{ds}{dt} &= -k_1 se + k_{-1} x_a \\ \frac{dp}{dt} &= k_c x_b - k_{-c} ep \\ \frac{dx_a}{dt} &= k_1 se - (k_{-1} + k_{ab}) x_a + k_{ba} x_b \end{aligned} \tag{6}$$

with conservation laws

$$\begin{aligned} x_b &= s_0 - (s + p + x_a) \\ e &= e_0 - x_a - x_b = e_0 - (s_0 - (s + p)) \end{aligned}$$

This model is able to fit the elastase kinetic data pretty well with physically meaningful parameters (Table S1), as shown in Figure S7. The best-fit solution reproduces exactly the behaviour obtained with eq. (11), reproduced in Figure 2 in the main text. Figure S8 shows the behaviour of the enzyme and intermediate complexes concentrations, as a function of time, for initial substrate concentration $s_0 = 750\ \mu M$.

##### 3.1.1 Quasi steady-state approximation

We notice that all kinetic steps in Table S1 are quite fast, with respect to our observation time range. Moreover, Figure S8 suggests that the intermediate complex $x_a$ reaches a roughly constant and low value in about 2000 $s$. We therefore try to impose $\frac{dx_a}{dt} = 0$ in (6) (quasi

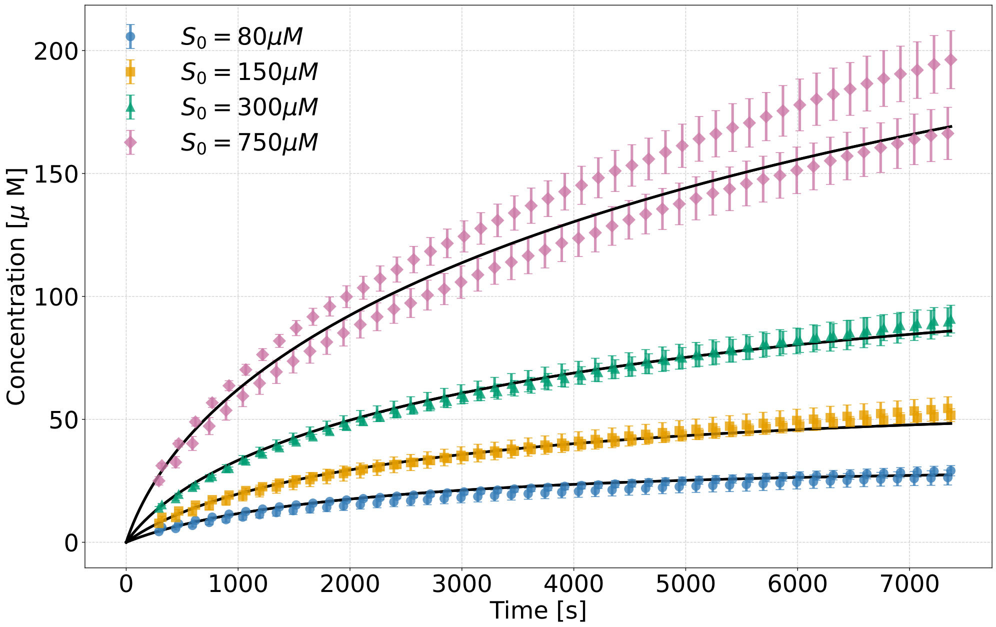


Figure S7: Three-step reversible mechanism with one product. Measured evolution of the product concentration in the absence of dextran (symbols) and best-fit with model (18) in the paper (solid lines), at different initial substrate concentrations. The data shown and the error bars represent averages and standard errors computed over 4 replicates.

Table S1: best-fit values for the three-step model.

| **Parameter** | **Value** |
|---|---|
| $X_R^2$ | 1.38 |
| $k_1$ | $(0.62 \pm 0.06) \times 10^3\ \mu M^{-1} s^{-1}$ |
| $k_{-1}$ | $(4.9 \pm 0.6) \times 10^6\ s^{-1}$ |
| $k_{ab}$ | $(42 \pm 6)\ s^{-1}$ |
| $k_{ba}$ | $(0.233 \pm 0.010)\ s^{-1}$ |
| $k_c$ | $(22 \pm 13)\ s^{-1}$ |
| $k_{-c}$ | $(0.9 \pm 0.5)\ \mu M^{-1} s^{-1}$ |

steady state approximation). Substituting the conservation law for $x_b$, this reads

$$k_1 se + k_{ba}(s_0 - (s + p + x_a)) - (k_{-1} + k_{ab})x_a = 0$$

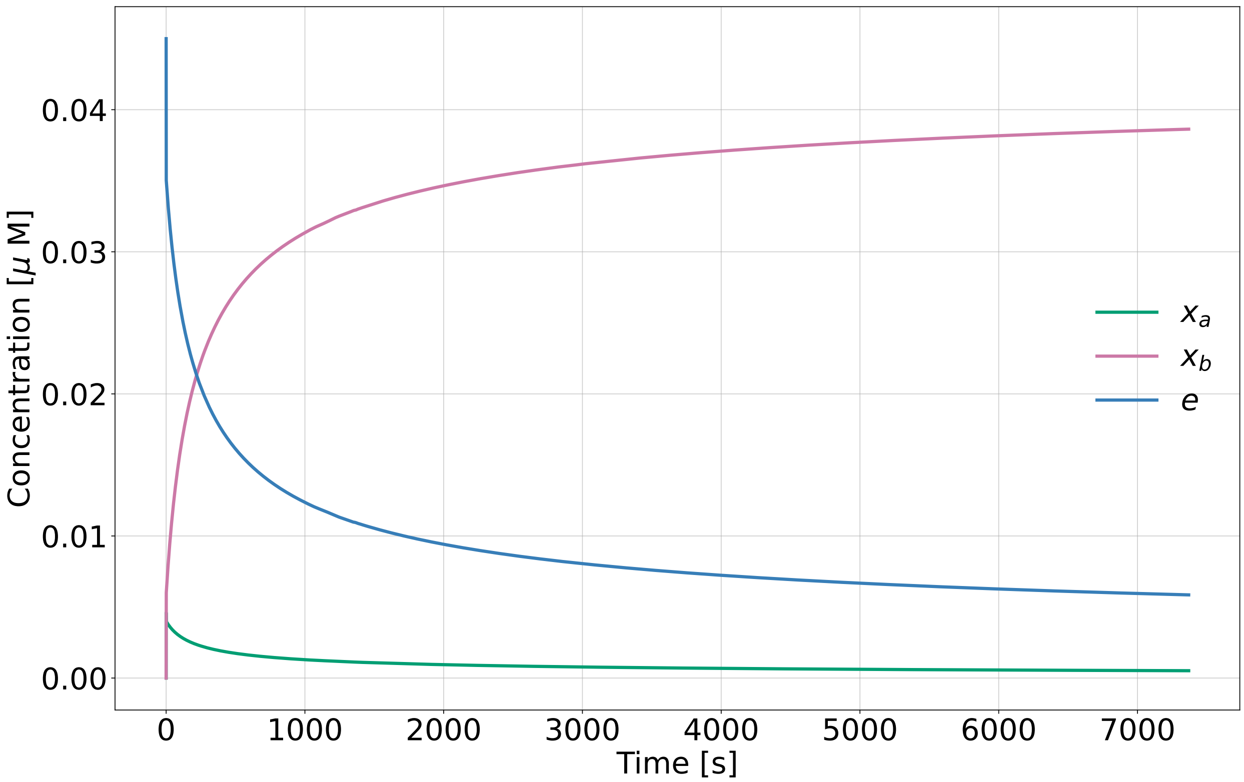


Figure S8: Free elastase and intermediate complexes concentration as a function of time, for $s_0 = 750$ $\mu$ M. Estimation of concentrations obtained integrating the system of ODEs (6) for the best-fit solution reported in Table S1.

This can be reorganized to find the $x_a$ value, choosing to express $s_0 - s + p$ as $e_0 - e$ (there are two different ways to write $x_a + x_b$ at any time following the conservation laws):

$$x_a = \frac{k_1 se + k_{ba}(e_0 - e)}{k_{-1} + k_{ab} + k_{ba}}$$

Then, substituting the expression for $x_a$ in (6), it reduces to a system of just two ODEs, one for the substrate and one for the product,

$$\frac{ds}{dt} = -k_1 se + k_{-1}\left(\frac{k_1 se + k_{ba}(e_0 - e)}{k_{-1} + k_{ab} + k_{ba}}\right)$$
$$\frac{dp}{dt} = k_c x_b - k_{-c} ep$$

The first eq. can be reorganized as follows:

$$\begin{aligned}\frac{ds}{dt} &= -k_1\left(1-\frac{k_{-1}}{k_{-1}+k_{ab}+k_{ba}}\right)se+\frac{k_{-1}k_{ba}}{k_{-1}+k_{ab}+k_{ba}}(e_0-e)\\ &= -k_1\left(\frac{k_{ab}+k_{ba}}{k_{-1}+k_{ab}+k_{ba}}\right)se+\frac{k_{-1}k_{ba}}{k_{-1}+k_{ab}+k_{ba}}(e_0-e)\end{aligned}$$

The system therefore takes the final form:

$$\begin{aligned}\frac{ds}{dt} &= -k_1\left(\frac{k_{ab}+k_{ba}}{k_{-1}+k_{ab}+k_{ba}}\right)se+\frac{k_{-1}k_{ba}}{k_{-1}+k_{ab}+k_{ba}}(e_0-e)\\ \frac{dp}{dt} &= k_c x_b - k_{-c}ep\end{aligned} \tag{7}$$

Then, Figure S8, suggests that $x_b \gg x_a$. In this case, $e_0 - e = x_a + x_b \simeq x_b$, and the system of equations (7) reduces exactly to the MM system (Eq. (12) in the main text), with $x_b \equiv x$ and:

$$\begin{aligned}k_{on} &= k_1\left(\frac{k_{ab}+k_{ba}}{k_{-1}+k_{ab}+k_{ba}}\right)\\ k_{off} &= \frac{k_{-1}k_{ba}}{k_{-1}+k_{ab}+k_{ba}}\end{aligned}$$

Using the values reported in Table S1, the estimation of the two effective kinetic constants is dominated by the significant difference between $k_1$ and $k_{ab}$ and $k_{ba}$. One finds:

$$\begin{aligned}k_{on} &\simeq k_1\frac{k_{ab}}{k_{-1}} = 0.53\times10^{-2}\ \mu M^{-1}s^{-1}\\ k_{off} &\simeq k_{ba} \simeq 0.233\ s^{-1}\end{aligned}$$

The estimations of $k_{on}$ and $k_{off}$ coincide exactly with the results of the fit reported in Table 3 in the main text, as well as those of $k_c$ and $k_{-c}$ that can be directly identified with the constants appearing in the MM model. The ratio $k_{ab}/k_{-1} \simeq 0.9\times10^{-5}$, shows that a conformational step $k_{ab}$, slow compared to the corresponding dissociation step $k_{-1}$, can be reabsorbed in a slow effective association constant $k_{on}$.

Figure S7 shows the fit obtained with the three-step mechanism, with one only product, that is presented in the appendix.

#### 3.2 Two products kinetic model

We report here a reversible model with two products, inspired by the serine protease mechanism, which can be defined in the following way:

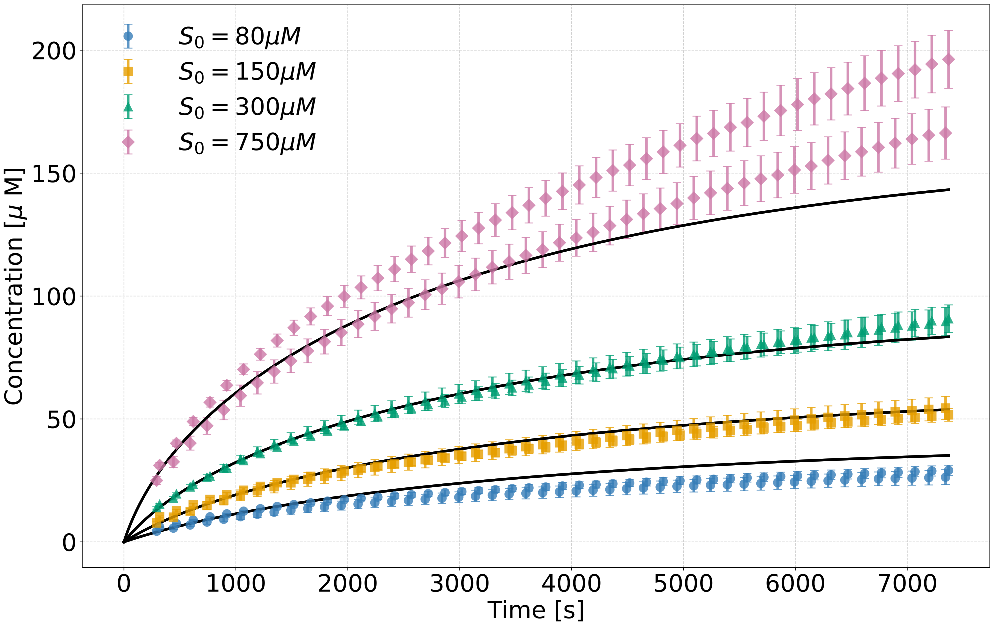


Figure S9: Three-step reversible mechanism with two products. Measured evolution of the product concentration in the absence of dextran (symbols) and best-fit with model (9) (solid lines), at different initial substrate concentrations. The data shown and the error bars represent averages and standard errors computed over 4 replicates.

$$\mathrm{S} + \mathrm{E} \underset{\mathrm{k_{off}}}{\overset{\mathrm{k_{on}}}{\rightleftharpoons}} \mathrm{X_1} \underset{\mathrm{k_{-2}}}{\overset{\mathrm{k_2}}{\rightleftharpoons}} \mathrm{X_2} + \mathrm{P_1} \tag{8}$$

$$\mathrm{X_2} \underset{\mathrm{k_{-3}}}{\overset{\mathrm{k_3}}{\rightleftharpoons}} \mathrm{E} + \mathrm{P_2}$$

It gives rise to the following system of ODEs for the concentrations:

$$\begin{aligned}
\frac{ds}{dt} &= -k_{on}se + k_{off}x_1 \\
\frac{dp_1}{dt} &= k_2x_1 - k_{-2}x_2p_1 \\
\frac{dp_2}{dt} &= k_3x_2 - k_{-3}ep_2
\end{aligned} \tag{9}$$

with conservation laws:

$$x_1 = s_0 - (s + p_1)$$

$$x_2 = p_1 - p_2$$

$$e = e_0 - x_1 - x_2$$

The best fit results are reported in Table S2 and the related solution is plotted in Figure S9. The quality of the fit is not as good as with models with one only product, as demonstrated by the higher $X_R^2$ and made apparent by direct comparison of the figures.

Table S2: Best-fit values for the three-step reversible mechanism with two products.

| **Parameter** | **Value** |
|---|---|
| $X_R^2$ | 4.66 |
| $k_{on}$ | $(0.73 \pm 0.08) \times 10^{-2}\ \mu M^{-1}s^{-1}$ |
| $k_{off}$ | $(1.6 \pm 0.5) \times 10^{3}\ s^{-1}$ |
| $k_2$ | $(2.3 \pm 0.5) \times 10^{3}\ s^{-1}$ |
| $k_{-2}$ | $(0.6 \pm 0.2) \times 10^{-2}\ s^{-1}$ |
| $k_3$ | $(1.1 \pm 0.4) \times 10^{3}\ s^{-1}$ |
| $k_{-3}$ | $(41 \pm 13)\ \mu M^{-1}s^{-1}$ |

### 4 Slow-homogeneisation model of kinetics in the presence of DEX450

In the main text we have developed a simple model that predicts a slow, quadratic initial time course for HNE kinetics in the presence of DEX450 solutions. In the slow-mixing hypothesis, the slow dynamics would reflect an additional rate-limiting phenomenon to be traced back to the properties of DEX450 solutions. These are likely heterogeneous at the microscopic scale, which might hamper substantially enzyme-substrate diffusive encounters at the start. The theoretical prediction for the early stages of the kinetics up to the onset of the linear MM regime reads

$$P(t, S_0) = P_0(S_0) + V(S_0)\left[t - \tau\left(1 - e^{-t/\tau}\right)\right] \simeq P_0(S_0) + \begin{cases} \frac{V(S_0)t^2}{2\tau} & \text{for } t \ll \tau \\ V(S_0)(t-\tau) & \text{for } t \gg \tau \end{cases} \tag{10}$$

Fig. S10 shows the results of simultaneous fits with a single relaxation time scale $\tau(c)$ at the considered concentrations of DEX450, $c$. The best-fit results of the slow time scale listed in Table S3 show that relaxation becomes increasingly slow as the crowding concentration increases, which confirm our set of hypotheses.

As mentioned in the main text, the free velocities $V(S_0, c)$ for the dataset at DEX450

Table S3: Primary free-$V(S_0, c)$ fit results. The quoted uncertainty on $\tau$ is the local Jacobian-based standard error and should be regarded as indicative only; it is not a formal confidence interval for this nonlinear, heteroscedastic problem.

| DEX450 (wt%) | $T_{\max}$ (s) | $\tau$ (s) | local SE($\tau$) (s) | RMSE ($\mu$M) | post-hoc $K_M$ ($\mu$M) |
|---|---|---|---|---|---|
| 5 | 1800 | 54 | 400 | 0.666 | 440 |
| 10 | 2000 | 334 | 13 | 0.363 | 427 |
| 15 | 2500 | 612 | 29 | 0.834 | 456 |
| 20 | 2500 | 971 | 55 | 1.019 | 784 |

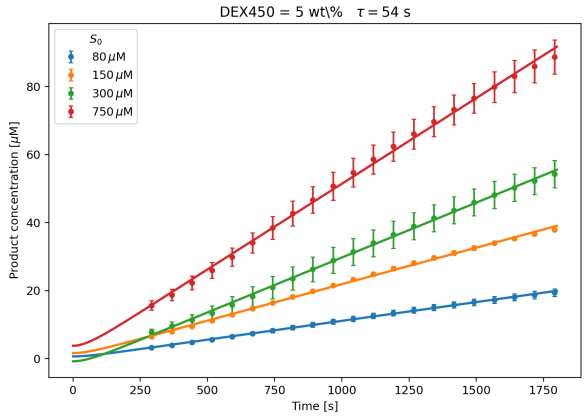


(a) 5 wt% DEX450.

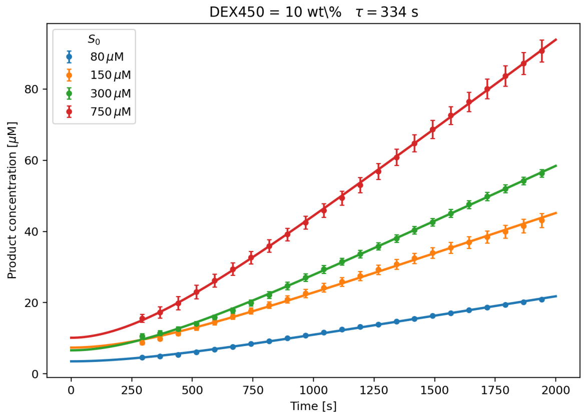


(b) 10 wt% DEX450.

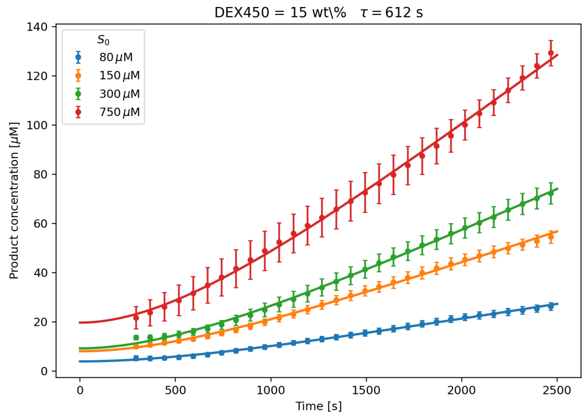


(c) 15 wt% DEX450.

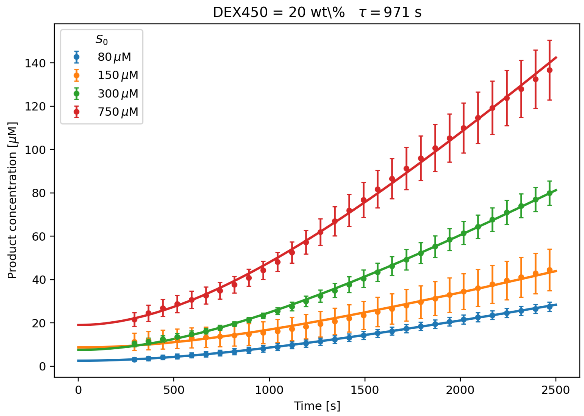


(d) 20 wt% DEX450.

Figure S10: Free$-V(S_0, c)$ global fits via Eq. (10) at different concentrations $c$ of DEX450.

concentration $c$, fitted individually, turn out to follow the expected MM trend, namely

$$V(S_0, c) = V_M(c) \frac{S_0}{S_0 + K_M(c)} \tag{11}$$

This is shown in Fig. S11.

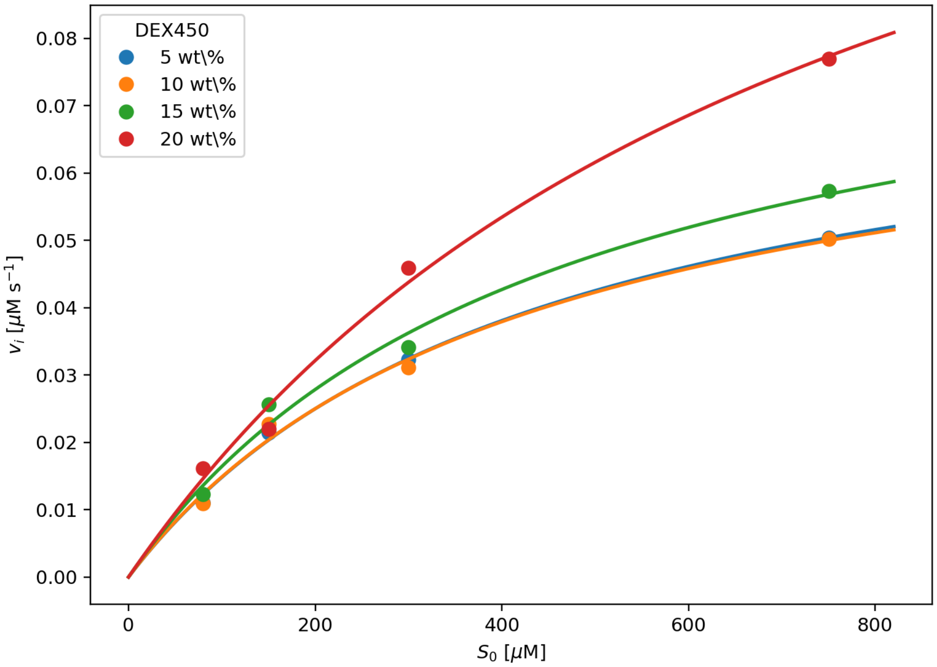


Figure S11: Free fitted asymptotic velocities $V(S_0, c)$ as a function of $S_0$. Solid lines are post-hoc Michaelis–Menten fits with Eq. (11) and were not imposed during the progress-curve fit. The apparent MM affinities are listed in Table S3.
